# Thalamic bursting and the role of timing and synchrony in thalamocortical signaling in the awake mouse

**DOI:** 10.1101/2021.07.09.451656

**Authors:** Peter Y Borden, Nathaniel C Wright, Arthur E Morrissette, Dieter Jaeger, Bilal Haider, Garrett B Stanley

**Affiliations:** Wallace H Coulter Department of Biomedical Engineering, Georgia Institute of Technology and Emory University, Atlanta, GA, USA; Emory University, Department of Biology, Atlanta, Georgia, USA

## Abstract

The thalamus controls transmission of sensory signals from periphery to cortex, ultimately shaping perception. Despite this significant role, dynamic thalamic gating and the consequences for downstream cortical sensory representations have not been well studied in the awake brain. We optogenetically modulated the ventro-posterior medial thalamus in the vibrissa pathway of the awake mouse, and measured spiking activity in the thalamus, and activity in primary somatosensory cortex (S1) using extracellular electrophysiology and genetically encoded voltage imaging. Thalamic hyperpolarization significantly enhanced thalamic sensory-evoked bursting, yet surprisingly the S1 cortical response was not amplified, but instead timing precision was significantly increased, spatial activation more focused, and there was an increased synchronization of cortical inhibitory neurons. A thalamocortical network model implicates the modulation of precise timing of feedforward thalamic population spiking, presenting a highly sensitive, timing-based gating of sensory signaling to cortex.

## INTRODUCTION

We explore the world through our sensory periphery, where sensors transduce the signals that ultimately give us perception of the world. The mammalian sensory thalamus gates information from the periphery to primary sensory cortices, controlling what signals do and do not make their way to cortex, thus playing a very critical role in sensing. Far from a static relay, the thalamus is under continuous influence by modulatory inputs from brainstem and feedback mechanisms from cortex (Sherman and Guillery, 2002; Sherman, 2005), most extensively explored in the context of sleep and epileptic seizures (Huguenard and McCormick, 2007; Fogerson and Huguenard, 2016). The large majority of detailed thalamic studies focusing on its potential role in sensory signaling have been either in ex-vivo/slice preparations or in intact/in-vivo preparations under anesthesia, where prominent thalamic properties such as synchronization of convergent thalamocortical projections and tonic/burst gating have been shown to strongly boost signaling under these conditions (Murray Sherman, 2001; Sherman, 2001; Swadlow and Gusev, 2001; Swadlow, 2002; Lesica and Stanley, 2004; Lesica et al., 2006; Wang et al., 2010; Stanley et al., 2012; Whitmire et al., 2016). Recent studies exploring the role of thalamus in the awake, unanesthetized brain have confirmed the critical role of thalamus in grossly modulating sensory cortex (Halassa et al., 2011; Poulet et al., 2012; Lewis et al., 2015; Reinhold et al., 2015; Yu et al., 2016; Wright et al., 2021). However, whether increases in thalamic excitability act to boost cortical sensory-evoked responses in the awake brain as predicted from ex-vivo and anesthetized studies is unknown.

Serving as the primary input to sensory cortex, properties of thalamic activity strongly shape the spontaneous, baseline activity of cortex, as well as the corresponding cortical response to ascending sensory inputs. It has been shown that even small changes in baseline membrane potential have appreciable effects on spontaneous firing of thalamic neurons (Béhuret et al., 2015), setting the overall tone of synaptic drive to cortex. The use of pharmacology to directly modulate thalamus (Godwin et al., 1996; Hirata and Castro-Alamancos, 2010; Poulet et al., 2012) or opto/microstimulation and pharmacology to indirectly affect thalamus through cortical (Olsen et al., 2012; Mease et al., 2014; Crandall et al., 2015; Li and Ebner, 2016; Kirchgessner et al., 2020) and subcortical (Aguilar and Castro-Alamancos, 2005) inputs has further causally revealed the extreme sensitivity of cortex to overall thalamic drive. One prominent characteristic of neurons in the thalamus is the presence of T-type calcium channels that are normally inactivated, but become de-inactivated through prolonged hyperpolarization (Jahnsen and Llinás, 1984a, 1984b; Sherman and Koch, 1986; Suzuki and Rogawski, 1989). Subsequent depolarizing inputs lead to calcium-mediated “bursts” of action potentials characterized by transient, high-frequency spiking, which is distinct from “tonic” spiking mediated through Na+/K+ channel dynamics. Most actively investigated in the context of sleep states and rhythmic discharge, the role of this mechanism in sensory signaling remains unclear, despite decades of speculation. At the synaptic level, thalamocortical high frequency bursting events have been shown to have a significant impact on downstream cortical activation (Swadlow and Gusev, 2001), where thalamic bursts are associated with an amplified post-synaptic response in recipient cortical neurons, thought to arise primarily from properties of the thalamocortical synapse. Coupled with sensitivity of cortex to the timing of thalamic inputs via the “window of opportunity” established by the disynaptic feedforward cortical inhibition (Pinto et al., 2000, 2003; Wehr and Zador, 2003; Wilent and Contreras, 2004, 2005; Gabernet et al., 2005; Isaacson and Scanziani, 2011), the gating of thalamic signaling through the aggregate effects of all these properties across the thalamic population is hypothesized to serve a critical role in processing sensory information (Crick, 1984; Pinto et al., 2000; Sherman and Guillery, 2002; Wilent and Contreras, 2004; Sherman, 2005). However, this has not been investigated extensively in the intact brain due to the lack of methodological approaches to precisely measure and manipulate thalamic properties. To date, no studies have utilized temporally precise, repeatable and reversible modulation to precisely measure the effects of thalamic burst/tonic gating in awake cortex at the level of population signals and single neuron firing.

Here, we directly determined how thalamic gating properties control sensory-evoked thalamic and cortical responses in the vibrissa pathway of the awake, head-fixed mouse. Instead of driving or silencing neural activity, optogenetic manipulation was used to modulate thalamus while recording extracellular thalamic and cortical activity and acquiring widefield cortical voltage imaging, using the voltage indicator ArcLight (Jin et al., 2012; Borden et al., 2017). We found that baseline thalamic firing rate was surprisingly invariant to optogenetic thalamic hyperpolarization through a transition from tonic to burst firing activity, and the baseline activity in S1 cortex was correspondingly invariant to thalamic hyperpolarization, following a brief transient increase in firing activity. In response to sensory stimuli, the thalamic hyperpolarization significantly enhanced the sensory-evoked thalamic bursting; however, the magnitude of the response in S1 cortex was not amplified, but instead slightly attenuated relative to baseline, despite the burst-dominated thalamic input. Notably, the sensory evoked response was also significantly more brief and spatially focused, and accompanied by an increase in the synchronization of the putative cortical Fast-spiking (FS) inhibitory neurons. A thalamocortical network model replicated these findings, and implicated changes in the precise timing and synchronization across the thalamic population as a likely mechanism underlying the experimental observations. Taken together, the results here point to timing rather than response magnitude as a fundamental feature of the thalamocortical circuit, presenting a dynamic, timing-based gating of sensory signaling to cortex.

## RESULTS

All experiments were conducted in the vibrissa pathway of the mouse (Figure 1A). To directly test the predictions of previous in-vitro and anesthetized work, we conducted the first experiment in the isoflurane-anesthetized mouse (illustrated in Figure 1B). In the lightly anesthetized mouse, we utilized extracellular electrodes to record whisker-evoked spiking activity in VPm thalamus, in the presence (LED, ∼17mW/mm^2^, 590nm) and absence (Control) of a hyperpolarizing optogenetic modulation of excitatory thalamic relay neurons expressing halorhodopsin (eNphR3.0, see Supplemental Figure S1), through a small fiber optic cable attached to the electrode (Figure 1B). Note that the inhibitory opsin was engaged with a constant illumination at a range of relatively low light levels, to induce sustained hyperpolarization, as opposed to complete inactivation. Note also that the total amount of light delivered here was in a range that has been previously shown to not induce significant heating of the surrounding tissue (see Methods), and control experiments indicate that there is no effect of the light alone in the absence of opsin expression (Supplemental Figure S4). Although the in-vivo optogenetic implementation precludes precise knowledge of the degree of hyperpolarization of the thalamic neurons due to variations in opsin expression, position of optic fiber relative to cells, etc., a separate set of in-vitro, brain slice experiments showed that VPm neurons were hyperpolarized by ∼15-25 mV for the light levels utilized (see Supplemental Figure S2). Given that in the in-vitro experiments, light was presented more directly to the VPm neurons expressing halorhodopsin, and that in the in-vivo experiments the presentation of light did not completely silence the neurons, it is likely that the in-vitro experiments were an upper-bound for the in-vivo case, and the actual hyperpolarization induced in-vivo was less than that of the in-vitro experiments.

**Figure 1.**
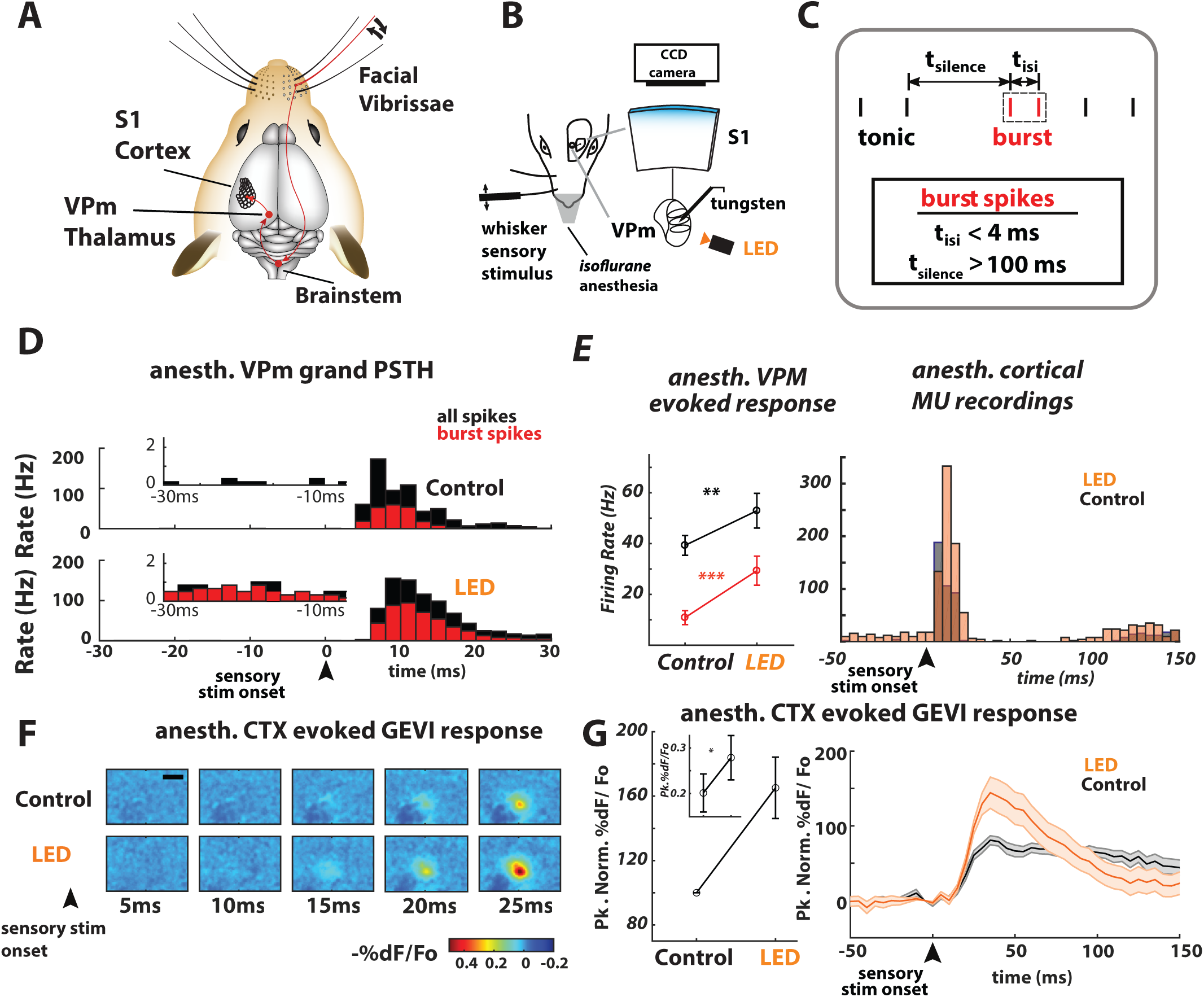
Thalamic hyperpolarization amplifies thalamic bursting and sensory-evoked cortical response in the anesthetized mouse. **A.** Pathway of the mouse vibrissa system from the facial vibrissae in the periphery, to brainstem, to thalamus, to S1. **B.** Experimental setup. Mice were injected with the viral vector eNphR3.0 (AAV5-CamIIKianse-eNphR3.0-mCherry) targeting the ventral posterior-medial (VPm) region of the thalamus and the viral vector ArcLight (AAV1-hysn1-ArcLightD- SV40) in the vibrissa region of the primary somatosensory cortex (S1). For optogenetic hyperpolarization, thalamic units were presented with constant illumination (LED, 590 nm, approx. 17 mW/mm^2^) from a 200µm optic fiber, with LED illumination starting at 0.5s preceding stimulus (t= -0.5s) and ending 0.5s after stimulus (t=0.5), while recording simultaneously with a single tungsten electrode. Note that unlike for the awake recordings, light presentation was not ramped. For cortical GEVI imaging, the entire cortical area was illuminated through the thinned skull at 465 nm with an LED and imaged with a CCD imaging setup (see Methods). **C.** Thalamic bursts were identified as two or more spikes with inter-spike interval (ISIs) less than 4ms, preceded by silence for 100ms or more. Thalamic spikes that were part of an identified burst were classified as putative burst spikes (red), and all else were classified as tonic spikes (black). **D.** Peri-stimulus time histogram (PSTH) for thalamic spiking in response to a single, punctate whisker stimulus (t=0) for the Control (no light, top panel) and thalamic hyperpolarized (LED, bottom panel) conditions, 29 units. Instantaneous firing rates (bin size 2 ms) shown for all spikes (black) and putative burst spikes (red). **E. Left** Mean sensory-evoked thalamic firing rate (n=29 thalamic units) over the 0 – 30 ms time window increased from the Control (no light) to the LED conditions, for all spikes (black, p= 0.0046, paired Wilcoxon signed-rank) and burst spikes (red, p= 4.5e-4, paired Wilcoxon signed-rank). Error bars represent mean +/- SEM. **Right**. Mean sensory-evoked Cortical multiunit (MU) firing rate from 102 trials (across 2 recordings, 1 mouse, 5ms bin size). **F.** Example session of GEVI imaging following the delivery of a punctate whisker stimulus at time t=0, for the Control (no light, top row) and LED (bottom row) conditions. Images are averaged across 51 trials. Black bar represents 1mm. **G. Left** - Mean normalized peak GEVI sensory-evoked response was larger for the LED as compared to the Control condition (n= 9 mice, 13 recording sessions). For this plot, animals and sessions were each normalized to their control levels, and the LED condition is reported relative to the control. **Left Inset** - raw relative peak evoked %dF/Fo, p=0.017, paired Wilcoxon signed-rank, n=13 recordings across 9 mice. **Right** - Time series of the normalized sensory-evoked GEVI signal, generated from the integrated fluorescence within the 0.2 x 0.2mm ROI in the Control (gray) and LED (orange) conditions. Error bars represent mean +/- SEM.

Neurons in VPm thalamus responded to a single, computer-controlled punctate whisker stimulation (1200 deg/sec, sawtooth waveform) with a brief, transient sequence of action potentials (Figure 1D, see Methods for single unit and multi-unit classification). Optogenetic thalamic hyperpolarization significantly increased the sensory-evoked thalamic bursting, as shown in Figure 1D (all spikes in black, putative T-type calcium “burst” spiking in red, classified according to the criterion in Figure 1C). Shown in the insets are the low baseline firing rates in the anesthetized condition. Overall, there was an increase in the evoked response from the Control to the LED condition (Figure 1E left, black, p=0.0046, paired Wilcoxon signed-rank, n=29 units), in large part driven by the increase in sensory evoked thalamic bursting (Figure 1E left, red, p=4.5e-4, paired Wilcoxon signed-rank, n=29 units). In a subset of experiments, we recorded the downstream cortical multiunit activity (Figure 1E right), and found a corresponding increase during the period of hyperpolarization (n=102 trials, 2 recordings, 1 mouse). Additionally, we simultaneously recorded cortical S1 activation at a meso-scopic scale with optical voltage imaging in primary somatosensory (barrel) cortex using a Genetically Engineered Voltage Indicator (GEVI, ArcLight, Borden et al., 2017) during optogenetic hyperpolarization of thalamus (see Methods). This approach enables the characterization of the overall spatiotemporal changes in S1 due to the thalamic manipulation (Supplemental Figure S4). The imaging was focused at a cortical depth consistent with layer 2/3 (see Methods). The fluorescence imaging of cortical S1 confirmed a corresponding amplification of the sensory evoked response during thalamic hyperpolarization (raw %dF/F_o_), as shown in an example session for the fluorescence images in Figure 1F and further quantified across the larger dataset in the left panel of Figure 1G (normalized relative peak evoked response, ∼60% increase, 1G inset raw relative peak evoked %dF/F_o_, p=0.017, paired Wilcoxon signed-rank, n=13 recordings across 9 mice). There was an amplification in the peak response, followed by a relatively fast transient decrease in fluorescence, followed by a gradual return to baseline over approximately ∼200ms. The results here using optogenetic manipulation of thalamus and wide-field voltage imaging of cortex are consistent with previous observations of spontaneous thalamic bursting and the impact on downstream synaptic targets in cortical layer 4 (Swadlow and Gusev, 2001). Importantly, the amplification in the peak response observed here under these conditions reflects what would have been expected from the observations in thalamus, if the cortical sensory representation were strongly by the feedforward thalamic bursting.

### Thalamic Hyperpolarization and Ongoing Thalamic and Cortical Activity in the Awake Brain

In a next set of experiments, we sought to characterize the influence of thalamic properties on gating of signaling to cortex in the awake mouse. Before turning to the sensory-evoked responses, we first probed the effect thalamic hyperpolarization has on baseline, ongoing activity that is significantly higher in the awake state as compared to under anesthesia. The baseline firing rate in the awake state is significantly higher than in the anesthetized condition, and we thus expect to see some effect of the optogenetic hyperpolarization on baseline firing. An example single-unit VPm recording (experimental setup in Figure 2A) is shown in the raster plot in Figure 2B. VPm neurons in awake mice exhibited moderate ongoing firing activity (4.9 Hz for this example). Upon presentation of a ramping optical input to thalamus (LED, 590nm, ∼17mW/mm^2^), the spiking activity underwent a qualitative change to a more sparse pattern of multiple action potentials, typical across all recorded cells. Bursts were present in the baseline (Control, no light) condition, but substantially more prevalent in the presence of the hyperpolarizing optogenetic input. This effect is summarized across all clearly identified single-units in Figure 2C (n=51 single VPm units), showing the average instantaneous firing rate for all spiking in black, and for the putative burst spiking in red (bands represent +/- 1 standard error of the mean, SEM). This qualitatively shows that following an initial decrease in firing rate at the onset of the light ramp, the VPm firing activity briefly rose above pre-stimulus baseline firing rate, before returning to a level near baseline. This was accompanied by an increase in the burst firing rate, suggesting that the return to pre-stimulus baseline firing rate was due to a compensatory transition from tonic to burst firing. Quantitatively, following the initial transient, the single-unit firing rate was relatively invariant from the Control (0 mW/mm^2^) to the hyperpolarized condition, with a slight increase for only the strongest of LED conditions (Figure 2D left, single-unit, Control vs LED p=0.2, Control vs Double LED p=0.02, unpaired two-sample t-test, n=54 SU recordings for Control, and n=51 for each of the LED conditions; ∼17mW/mm^2^ - LED or ∼35mW/mm^2^ - Double LED). Note that the firing rate in the LED conditions was calculated from the 250-750ms window, after the initial transient decrease in firing rate. There was a corresponding significant increase in the firing rate associated with bursting (red, Control vs LED p=3e-13, Control vs Double LED p= 1e-15, unpaired t-test), indicating a compensatory shift from tonic to burst firing with the thalamic hyperpolarization resulted in a significant increase in the burst ratio metric from the Control to hyperpolarized condition (Fig. 2D, right, Control vs LED p=9e-13, Control vs Double LED p=1.8e-11, Wilcoxon rank-sum test). Further examination showed that increasing hyperpolarization also increased the number of spikes per burst (illustrated in left of Figure 2E), which partially offsets the loss of tonic spiking (right of Figure 2E, Control vs LED p=2.3e-5, Control vs Double LED p=1.7e-6, Wilcoxon rank-sum test, n=54 for the Control and n=51 for both LED conditions).

**Figure 2.**
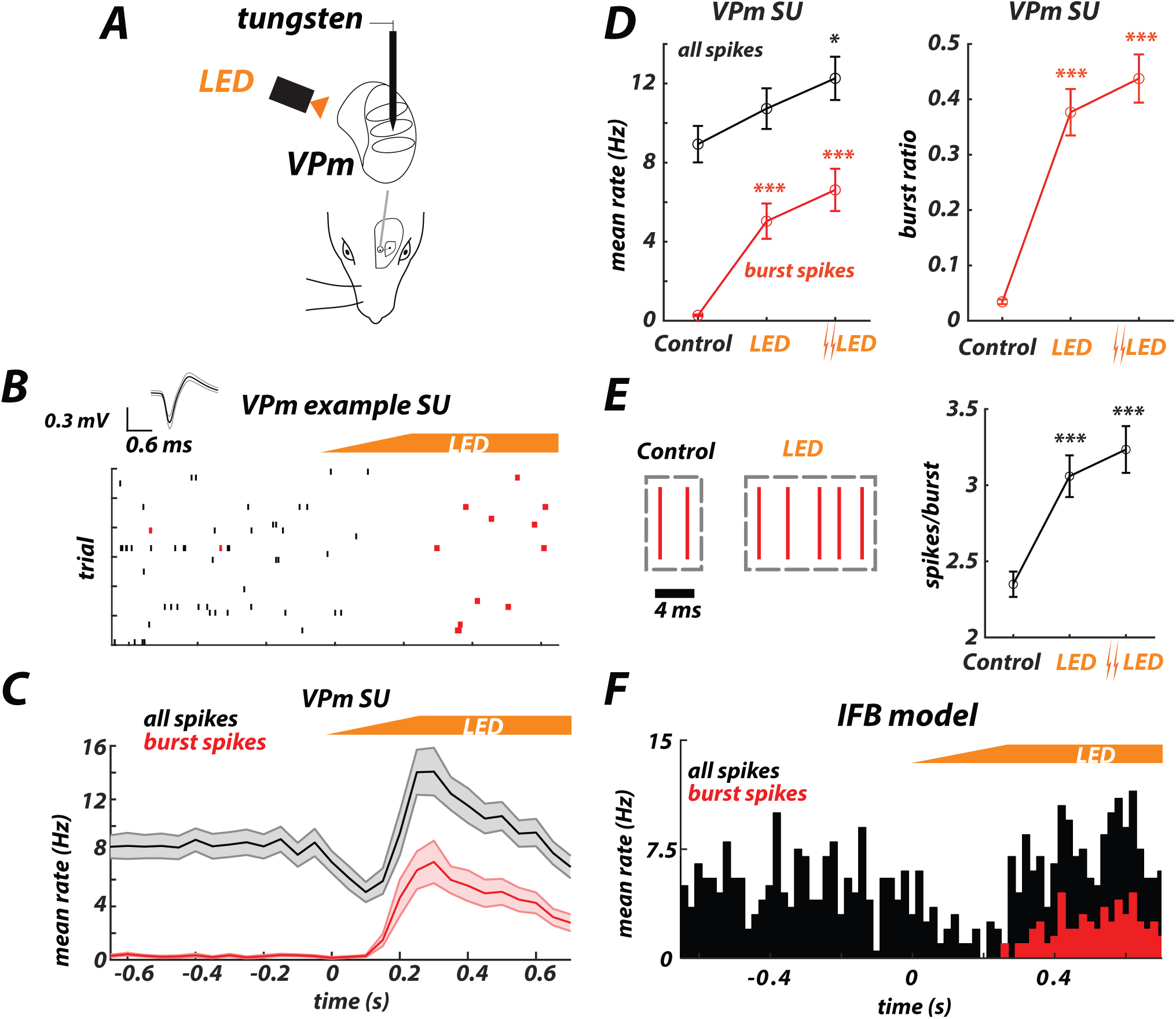
Thalamic baseline firing rate is invariant to optogenetic hyperpolarization via a tonic/burst switch. **A.** Experimental setup. Mice were injected with the viral vector eNphR3.0 (AAV5-CamIIKianse-eNphR3.0-mCherry) targeting the ventral posterior-medial (VPm) region of the thalamus. On each trial, thalamic units were presented with light for 1.5 s (590 nm, LED approx. 17mW/mm^2^ ramp, Double LED approx. 35mW/mm^2^ ramp – see Methods) from a 200µm optic fiber and recorded simultaneously with a single tungsten electrode. **B.** Example extracellular single trial rasters depicting effects of thalamic hyperpolarization on spiking in VPm thalamus. Black indicates tonic spikes; red indicates burst spikes. Recorded spike waveform shown at top. **C.** Aggregate single-unit (SU, n=51 units) PSTH for all spikes (black) and burst spikes (red). Apparent is an initial decrease in overall firing rate following presentation of light, followed by a return to the pre-hyperpolarization level. Bands represent are mean +/- SEM. **D. Left -** Mean single-unit thalamic firing rate over the 250-750ms time window from the Control (no light) to the LED condition, for all spikes (black, Control vs LED p=0.2, Control vs Double LED p=0.02, unpaired two-sample t-test, n=54 for Control, and n=51 for both LED conditions) and burst spikes (red, Control vs LED p=3e-13, Control vs Double LED p=1e-15, unpaired t-test). **Right -** Increase in burst ratio from the Control to LED condition, where burst ratio is defined as the number of burst spikes divided by the total number of spikes (Control vs LED p=9e-13, Control vs Double LED p=1.8e-11, unpaired Wilcoxon rank sum test). **E. Left -** Example bursts for the Control and LED conditions. **Right -** Increase in the mean number of spikes per burst from the Control to the LED condition (Control vs LED p=2.3e-5, Control vs Double LED p=1.7e-6, unpaired Wilcoxon rank sum test, n=54 for Control and n=51 for each of the LED conditions). Error bars represent mean +/- SEM. **F.** Simulated spontaneous activity from Integrate-and-fire-or-burst (IFB) model with hyperpolarizing input at time t=0, showing all spikes (black) and burst spikes (red). See Methods and Supplemental Figure S3.

Although it is not surprising that the thalamic burst mechanism was engaged by the optogenetic hyperpolarization, it is surprising that the net baseline firing rate was not decreased, but was instead relatively invariant to the thalamic hyperpolarization. This effect was not as apparent in the anesthetized condition when the baseline thalamic firing rate is near zero, but emerges in the awake state where baseline thalamic firing rate is significantly higher. To investigate possible causes, we constructed an integrate-and-fire or burst (IFB) model of thalamic firing that has been utilized in previous studies (Smith et al., 2000; Lesica and Stanley, 2004; Lesica et al., 2006), where the resting membrane potential was set by the level of activation of the hyperpolarizing opsin (see Methods). When the model neuron was set to have a spontaneous, baseline firing rate consistent with our observations (∼5 Hz), the effect of the hyperpolarizing opsin activation indeed replicated this phenomenon (Figure 2F). An initial decrease in firing rate at the onset of the light ramp was followed by a return to pre-light baseline level of firing (compare with Figure 2C). Moreover, as in the experimental observation, the return to the baseline firing rate in the hyperpolarized condition is dominated by the increase in bursting activity in the IFB model (Figure 2F, red). The exact combination of the baseline activity and intensity of the hyperpolarizing input strongly influenced the net resultant effect (Supplemental Figure S3). Prolonged periods of halorhodopsin activation have been shown to potentially alter the reversal potential of chloride (Raimondo et al., 2012). However, due to the short timescales of LED stimulation used in this work (1-1.5s), and the large transition to burst firing, the effect shown here is more likely driven by low voltage activation of T-type calcium channels, further reinforced by the IFB model.

To uncover the downstream effects of the above observations in VPm, we performed the same optogenetic manipulations during multi-electrode recording of single-unit activity in S1 in the awake mouse (Figure 3A). Note that for this analysis, while these S1 recordings were combined across cortical layer and cell type, subsequent analyses demonstrate similar trends across cell types and for targeted layer 4 recordings (see Figure 7 and Supplemental Figure 5). Shown in the top panel of Figure 3B is a raster plot from one example session (10 trials each from 15 simultaneously recorded units), in response to the optogenetic hyperpolarization of thalamus with the LED (again 590nm, 17mW/mm^2^). This example demonstrates a clear transient increase in spiking with the thalamic hyperpolarization, typical of the cortical recordings. The bottom panel of Figure 3B shows the aggregate single-unit firing rate, with a clear transient increase in firing rate at the light onset (bands represent +/- 1 SEM), with a gradual return to the baseline, pre-hyperpolarization level of firing rate at steady-state. Figure 3C summarizes this trend, showing the mean single-unit firing rate across recorded cortical single-units in the Control condition (0mW/mm^2^) versus activation of halorhodopsin in thalamus (LED - 17mW/mm^2^ or Double LED - 35mW/mm^2^), for the initial increase in firing rate during a transient period (300-350 ms, open symbols) and the subsequent return to baseline firing rate at steady state (700-750 ms, closed symbols). There was a significant transient increase in the cortical firing rate across both led conditions (transient, Control vs LED p=5.4e-14, Control vs Double LED p= 1.4e-15, Wilcoxon signed-rank test, n=118 units). After the transient increase, we observed a subsequent return to near baseline, either with no net effect or a marginal increase in the rate (steady-state, Control vs LED p=0.2, Control vs Double LED p= 0.013, Wilcoxon signed-rank test). Interestingly, following thalamic hyperpolarization, the cortical firing rate appeared to begin its initial transient increase while the thalamic activity remained relatively low (Figure 2), which could reflect an increased sensitivity of the thalamocortical synapse following the initial period of quiescence (Swadlow and Gusev, 2001; Swadlow et al., 2005).

**Figure 3.**
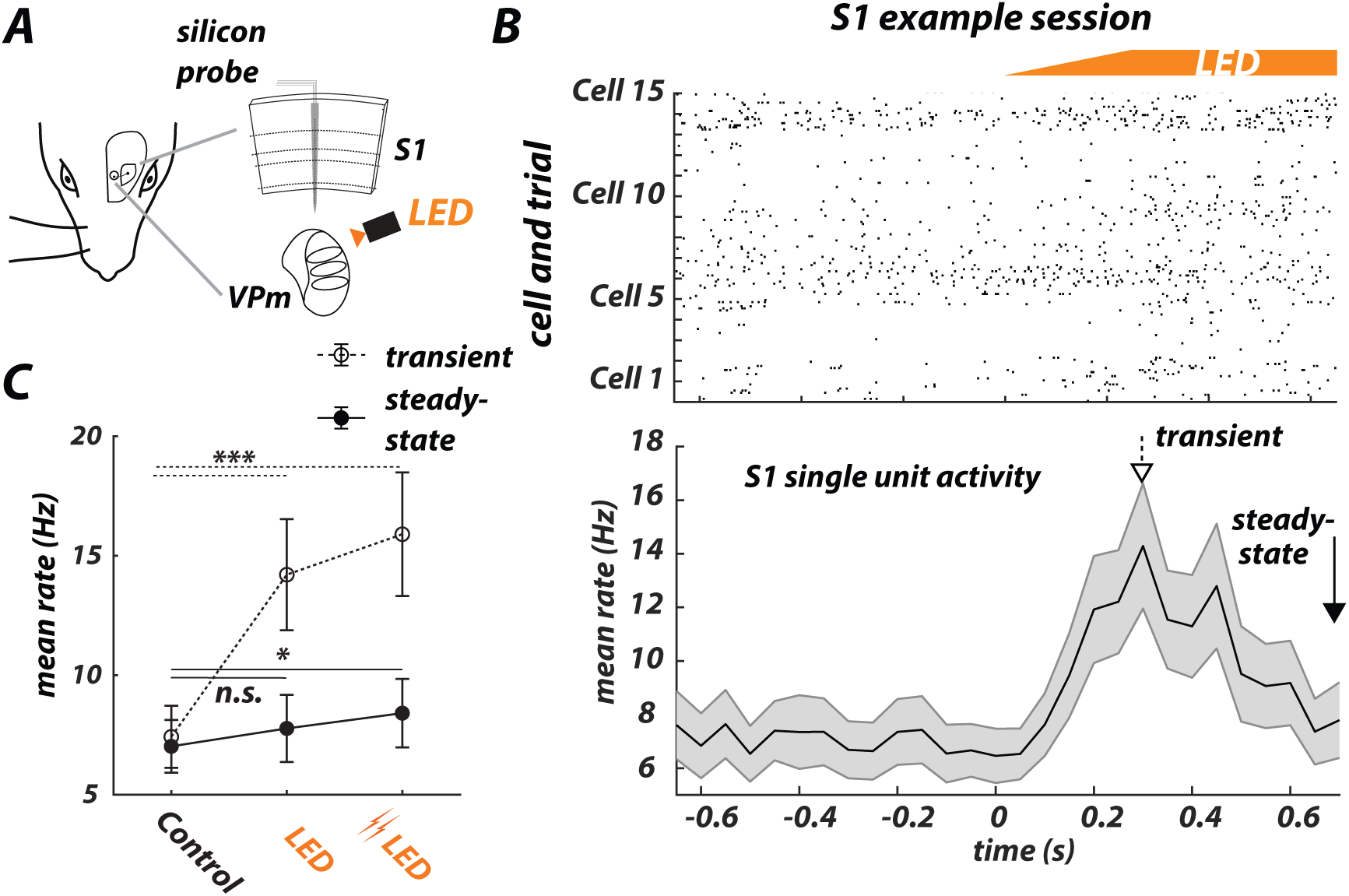
Cortical S1 baseline firing rate is invariant to optogenetic hyperpolarization of thalamus after transient increase. **A.** Experimental setup. Mice were injected with the viral vector eNphR3.0 (AAV5-CamIIKianse-eNphR3.0-mCherry) targeting the ventral posterior-medial (VPm) region of the thalamus. Thalamic units were continuously presented with light (590 nm, LED approx. 17mW/mm^2^ ramp, Double LED approx. 35mW/mm^2^ ramp) from a 200µm optic fiber, while cortical single-units were recorded simultaneously with a laminar multi-electrode within an identified cortical column (barrel) – see Methods. **B. Top -** Example extracellular rasters of cortical activity, depicting effects of thalamic hyperpolarization on cortical spiking, across trials and simultaneously recorded units (10 trials each from 15 units). **Bottom -** Aggregate PSTH across all recorded trials and cortical single-units (n=118). Bands represent mean +/- SEM. Highlighted are the transient and steady-state portions of the cortical response to the light. **C.** Mean cortical firing rate in the Control and LED conditions, showing an increase in the transient (open symbol, 300-350ms, Control vs LED p=5.4e-14, Control vs Double LED p=1.4e-15, Wilcoxon signed-rank test) and a return to near steady-state (closed symbol, 700-750ms, Control vs LED p=0.2, Control vs Double LED p=0.013, Wilcoxon signed-rank test) portions of the response. Error bars represent mean +/- SEM.

Taken together, the results here suggest that in response to the optogenetically induced thalamic hyperpolarization, the thalamic VPm neurons in the awake mouse exhibit a transient decrease in baseline firing rate, followed by a return to approximately pre-hyperpolarization level that is at least partially explained by a trade-off between tonic and burst firing, while the downstream activity in cortical S1 exhibits a transient increase in baseline firing rate, followed by an eventual return to approximately the original baseline rates.

### Thalamic Hyperpolarization and Sensory Evoked Thalamic and Cortical Activity

We next recorded the sensory evoked single-unit activity in VPm thalamus in the presence and absence of thalamic hyperpolarization in the awake mouse (Fig. 4A). Neurons in VPm thalamus responded to a single, computer-controlled punctate whisker stimulation (1200 deg/sec, sawtooth waveform) with a brief, transient increase in single-unit activity, which was significantly reshaped by thalamic hyperpolarization (Fig. 4B; PSTH of all spikes (black) across single-units, 2 ms bins, LED approximately 17mW/mm^2^, n=54 VPm single units for control, n=51 for LED). Also shown are the PSTHs comprised of putative burst spikes only (red), present in the Control condition but appearing to be significantly increased in the LED condition. To further quantify this effect, we tested a range of LED intensities to determine the impact of various levels of thalamic hyperpolarization on the evoked response (Figure 4C; LED approximately 17mW/mm^2^, and Double LED approximately 35 mW/mm^2^). Overall, there was a significant increase in sensory-evoked VPm bursting with thalamic hyperpolarization (Figure 4C, left, burst spikes (red), Control vs LED p=0.003, Control vs Double LED p=1.6e-4, unpaired Wilcoxon rank sum, n=54 VPm single-units for Control and n=51 for both LED conditions). However, for the same set of recordings there was a surprisingly invariant overall sensory-evoked response magnitude across VPm single units (Figure 4C, left, all spikes (black), Control vs LED p=0.9, Control vs Double LED p=0.4, unpaired two sample t-test). There was a corresponding significant increase in single-unit burst ratio (Figure 4C, right, Control vs LED p=1.3e-5, Control vs Double LED p=2.4e-6, Wilcoxon rank-sum, n=54 VPm single-units for Control and n=51 VPm single-units for both LED conditions). Taken together, single-unit VPm analyses reveal a boosting of the sensory evoked thalamic bursting with thalamic hyperpolarization, yet a surprisingly invariant overall mean evoked rate.

**Figure 4.**
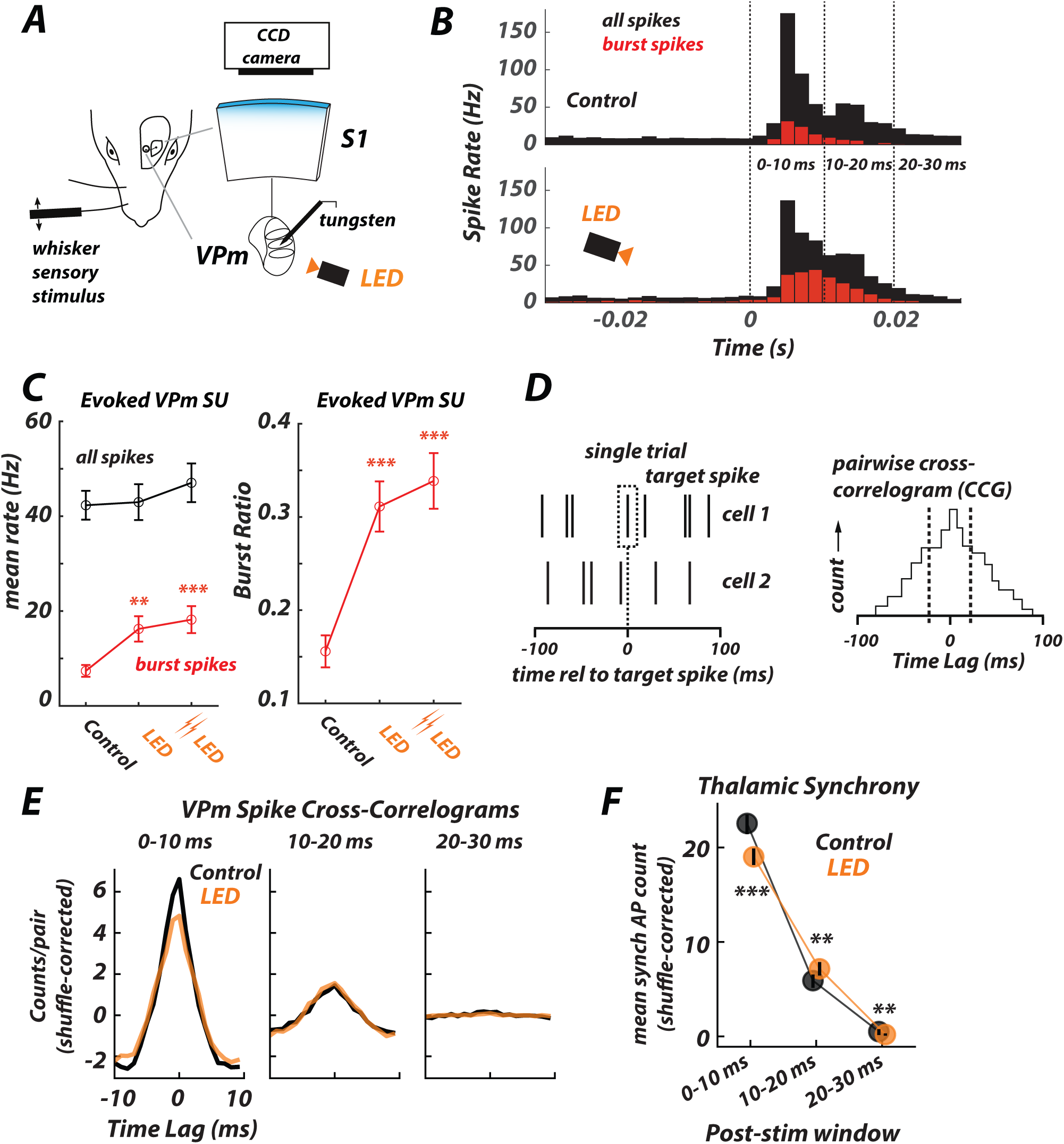
Thalamic hyperpolarization enhances the thalamic sensory-evoked bursting response in the awake mouse. **A.** Experimental setup. Mice were injected with the viral vector eNphR3.0 (AAV5-CamIIKianse-eNphR3.0-mCherry) targeting the ventral posterior-medial (VPm) region of the thalamus and the viral vector ArcLight (AAV1-hysn1- ArcLightD- SV40) in the vibrissa region of the primary somatosensory cortex (S1). For optogenetic hyperpolarization, on each trial thalamic units were presented with light for 1.5 s (590 nm, LED 29approx. 17mW/mm^2^ ramp, Double LED 29approx. 35mW/mm^2^ ramp – see Methods) from a 200µm optic fiber, with LED illumination beginning at 0.75s preceding stimulus delivery, and continuing for 0.75s after stimulus delivery, while recording simultaneously with a single tungsten electrode. For cortical GEVI imaging, the entire cortical area was illuminated through the thinned skull at 465 nm with an LED and imaged with a CCD imaging setup (see Methods). **B.** Grand single unit PSTH for thalamic spiking in response to a single, punctate whisker stimulus (t=0) for the Control (no light, top panel, n=54 units) and LED (bottom panel, n=51 units) conditions. Instantaneous firing rates (bin size 2 ms) shown for all spikes (black) and putative burst spikes (red). **C. Left** – Mean sensory- evoked thalamic response over 0-30ms time window for single-unit VPm recordings, for all spikes (black, Control vs LED p=0.9, Control vs Double LED p=0.4, unpaired two-sample t-test, n=54 for Control, and n=51 for both LED conditions), and for burst spikes (red, Control vs LED p=0.003, Control vs Double LED p=1.6e-4, unpaired Wilcoxon rank sum, n=54 for Control, n=51 for both LED conditions). Error bars represent mean +/- SEM. **Right** – Mean single-unit sensory-evoked thalamic burst ratio over the 0–30 ms time window (Control vs LED p=1.3e-5, Control vs Double LED p=2.4e-6, unpaired Wilcoxon rank sum, n=54 for Control, and n=51 for both LED conditions). **D.** Construction of spike cross-correlogram (CCG) from pair-wise VPm spiking. **E.** Grand mean CCGs of the sensory-evoked response for the Control and LED conditions, when restricting spiking data to the [0-10ms], [10-20ms], and [20-30ms] post-stimulus time windows (n=187 simultaneously recorded pairs from 46 VPm single units, across 6 recording sessions). **F.** Mean shuffle-corrected synchronous spike count for the [0-10ms], [10-20ms], and [20-30ms] post- stimulus time windows. All reported values are mean synch AP count +/- 99.95% confidence interval, resampling relative spike times with replacement ([0-10 ms], Control to LED, p < 0.001; [10-20 ms], p < 0.01; [20-30 ms], p < 0.01).

Note that upon more careful inspection of the VPm PSTHs, thalamic hyperpolarization results in a qualitative decrease in the sensory-evoked response in the first 10 ms, and a subsequent increase in the sensory-evoked response at later times. This could indicate changes in synchronous thalamic firing over the course of the sensory response, which is a critical factor in determining the downstream cortical response. To explicitly quantify this, we analyzed a large subset of the VPm recordings that were obtained from multi-electrode arrays. The multi-electrode arrays yielded a portion of simultaneously recorded VPm single-units, thus enabling us to analyze pair-wise properties. Figure 4D illustrates the formation of the spike cross-correlogram (CCG) from the spiking activity of a simultaneously recorded pair of neurons (see Methods). We applied this analysis across pairs of simultaneously recorded VPm pairs, restricting the spiking activity to three non-overlapping windows following the whisker stimulus (0-10ms, 10-20ms, and 20-30ms). The resulting grand mean CCGs for the Control and LED condition are shown in Figure 4E. Qualitatively we observe a decrease in the central peak of the CCG for the earliest post-stimulus time window (0-10ms, left) from the Control to the LED condition, and very little change for the other two time windows (10-20ms middle, 20-30ms right). We quantified this observation by computing the area under the CCG in the central window [- 5ms, +5ms], representing the amount of synchronous spiking across the pairs. Most notably, the LED induced a decrease in the synchronous firing for the earliest post-stimulus time window (0 – 10 ms), when synchronous spike rates were generally highest (Control vs. LED, p < 1e-3, n=187 pairs from 46 VPm units, comparison of confidence intervals from re-sampling with replacement, see Methods). Synchronous spike rates also changed in the two later response windows, when synchronous rates were generally lower, with a slight increase for the (10-20 ms) window and a slight decrease for the (20-30 ms) window (10 – 20 ms: p < 0.01; 20 – 30 ms: p < 0.01, n=187 pairs from 46 VPm units). Thus, taken together, the VPm data reveal a significant boosting of sensory-evoked burst firing with thalamic hyperpolarization, accompanied by a relatively invariant overall sensory-evoked response magnitude, and a surprising decrease in the synchrony of the prominent early thalamic response.

To examine the downstream consequences of the observed changes in the sensory-evoked response in thalamus, we recorded cortical S1 activation with wide-field optical voltage imaging during optogenetic hyperpolarization of thalamus in the awake mouse (illustrated in Figure 5A, see Methods). Note further that fluorescent imaging approaches like this are differential in nature (responses are based on relative not absolute changes in fluorescence), and are thus not well suited to capture overall absolute levels of ongoing background activity, but instead are targeting evoked responses relative to background. Based on previous findings (Swadlow and Gusev, 2001), the results in the anesthetized mouse, and the significant boosting of sensory-evoked bursting in thalamus with thalamic hyperpolarization in the awake mouse, we expected to see a corresponding amplification of the cortical sensory-evoked response. Shown in Figure 5A is a trial averaged sequence of the sensory-evoked fluorescence images. Following the deflection at time 0, a localized change in fluorescence began to emerge 10-15 ms later. Surprisingly, thalamic hyperpolarization did not result in an increased cortical response, but instead induced an apparent modest decrease in sensory evoked S1 activity. As quantified in Figure 5B, the peak fluorescence showed an approximately 20-40% decrease with this level of thalamic hyperpolarization (Control vs LED p=0.020, Control vs Double LED p=0.0039, paired Wilcoxon sign rank test, 2 mice over 9 sessions for LED conditions). Note that the right panel of Figure 5B is the same data as in the left panel, with each dataset normalized to the Control condition, to show a percent decrease in the peak fluorescence.

**Figure 5.**
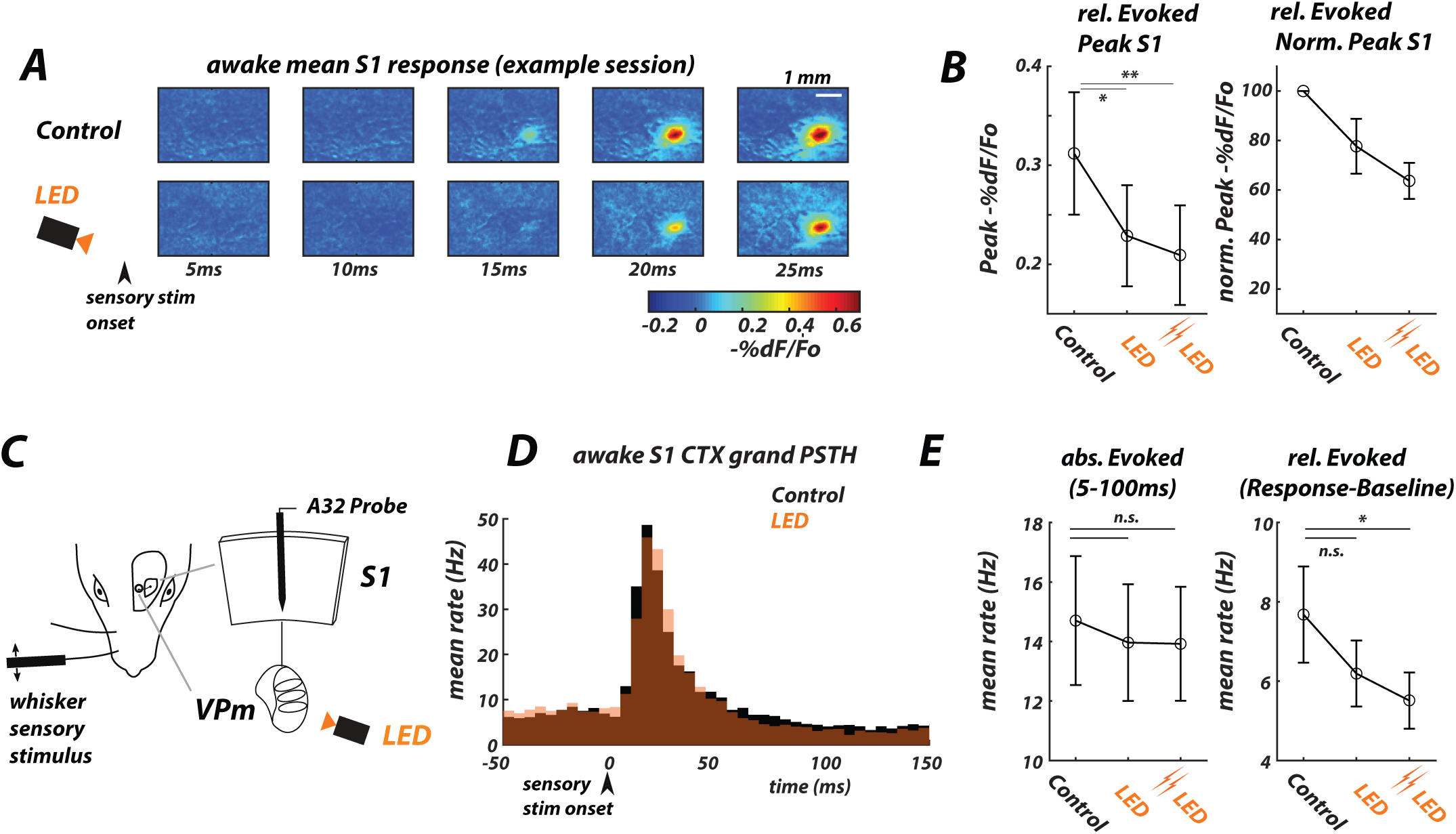
Cortical sensory-evoked response magnitude is invariant to thalamic hyperpolarization in the awake mouse. **A.** Example session of GEVI imaging following the delivery of a punctate whisker stimulus at time t=0, for the Control (no light, top row) and LED (bottom row) conditions. Images are averaged across 51 trials. **B. Left** – Peak GEVI sensory- evoked response (between 0-110ms post stimulus) slightly decreased from the Control to the LED condition (Control vs LED p=0.020, Control vs Double LED p=0.0039, paired Wilcoxon signrank, n=9 recordings from 2 mice). **Right –** Mean normalized peak GEVI sensory-evoked response for the Control and LED conditions (n=9 recordings). Before combining, animals and sessions were each normalized to their Control levels, and the LED condition is reported relative to the Control. **C.** In separate experiments, we hyperpolarized the thalamus while simultaneously recording the S1 Cortical response using a multichannel electrode. **D.** Cortical Grand PSTH evoked sensory response (t=0) across all recorded single units (n=118 units, 5ms bins) for Control (black) and LED (orange) conditions. **E. Left** Average absolute evoked cortical response remained approximately invariant during LED on conditions across all cortical recorded units (n=118). Evoked sensory response period defined as between 5-100ms post stimulus. **Right.** Relative evoked cortical response decreased with increasing LED intensities (Control vs LED p=0.20, Control vs Double LED p=0.034, paired Wilcoxon signrank, n=118 units). Relative evoked response defined as the absolute response minus the preceding baseline activity −50-0ms pre-stimulus. Error bars represent mean +/- SEM.

Finally, in a separate set of experiments, we hyperpolarized thalamus while simultaneously recording the S1 Cortical SU responses using a multichannel electrode (illustrated in Figure 5C). The corresponding PSTHs for responses to a punctate whisker stimulus (1200 deg/sec, sawtooth waveform) are shown in Figure 5D (Control (0mW/mm^2^) vs. LED (17mW/mm^2^) conditions). The average absolute evoked cortical response remained approximately invariant during LED conditions across all cortical recorded units (Figure 5E, left, 5 to 100ms post-stimulus, n=118). However, the evoked cortical spiking relative to baseline decreased with increasing LED intensities (Control vs LED p=0.20, Control vs Double LED p=0.034, paired Wilcoxon signrank, n=118 units, across 3 mice), consistent with the GEVI recordings (Figure 5B), which are relative measures by construction. Note that the recorded cortical activity here was aggregated across cell type and cortical layer, but subsequent analyses will address cell type and layer (see Figure 6 and Supplemental Figure S5). Importantly, similar trends were observed through analysis restricted to putative layer 4 cortical neurons (Supplemental Figure S5A and B).

**Figure 6.**
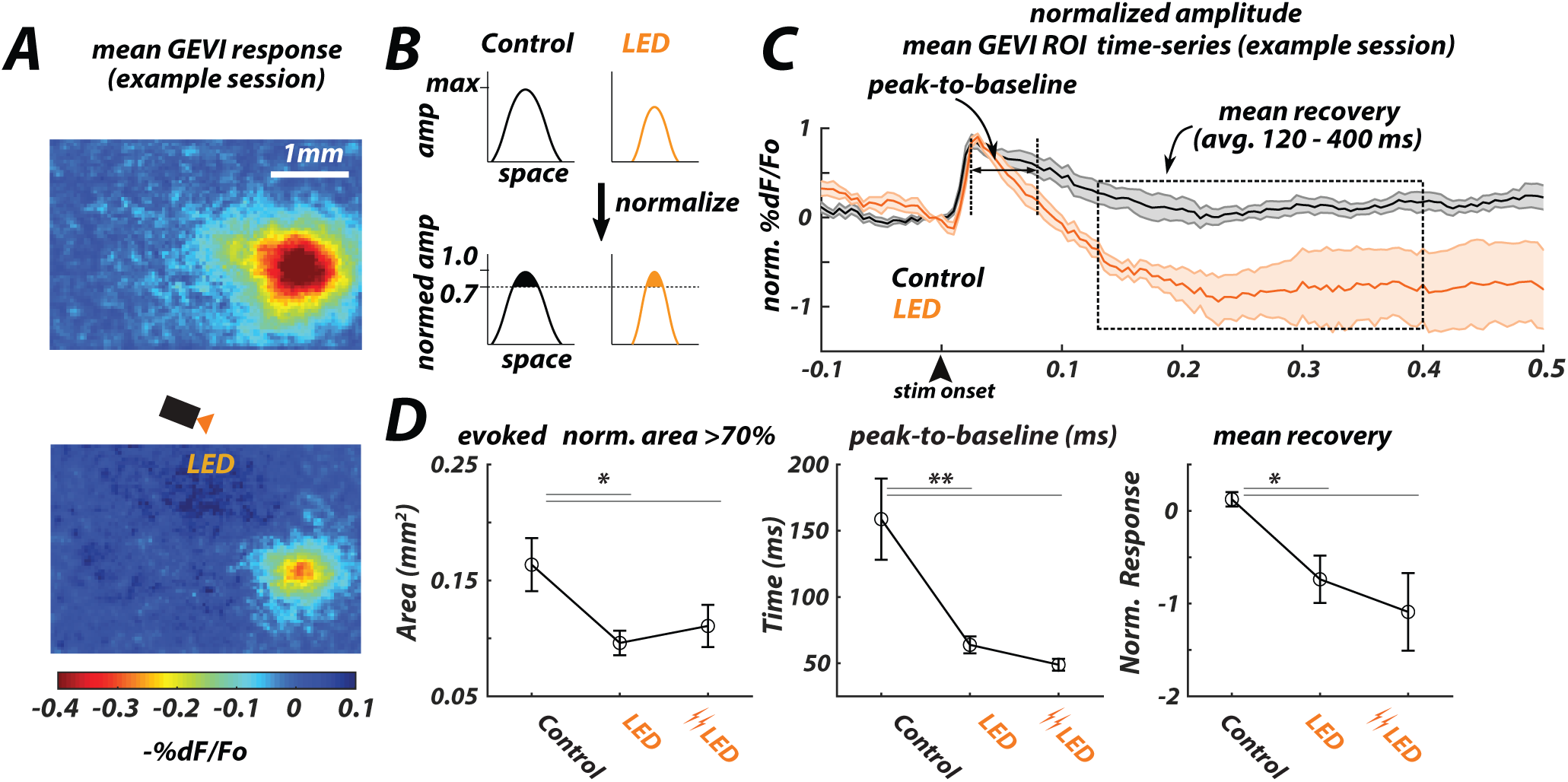
Thalamic hyperpolarization results in a spatially sharpened and temporally narrowed cortical sensory evoked response. **A.** GEVI imaging frames at peak response for the control (top) and LED (bottom) conditions for an example session. **B.** Cartoon illustration of the coupling between amplitude and spatial area in cortical imaging. **Top -** A reduction in evoked fluorescence amplitude can be qualitatively perceived as a reduction in spatial area of activation, due to the “iceberg” effect. **Bottom -** However, the spatial area of activation above a fixed threshold after normalization to the peak reveals the true effects on the area. **C.** Normalized amplitude time series of fluorescence within R0I for Control (black) and thalamic hyperpolarized (orange, LED), n= 9 recordings from 2 mice) conditions. Bands represent mean +/- SEM. **D.** Summary analyses (Error bars represent mean +/- SEM). Note, for this figure all conditions are normalized to their respective peak responses, as portrayed in B. **Left** - Mean spatial activation area of the normalized sensory-evoked response decreased from the Control to the LED condition (Control vs LED p=0.025, Control vs Double LED p=0.046, paired t-test, n= 9 recordings). **Middle** - The peak- to-baseline (defined in Figure 6C), calculated as the mean time from the peak of the fluorescence to the first return to pre-stimulus baseline (ms), decreased from the Control to the LED condition (Control vs LED p=0.0059, Control vs Double LED p=0.0019, unpaired t-test, n= 8 Control recordings, 9 LED recordings, note one of the Control recordings did not return to baseline). **Right** – The recovery (defined in Figure 6C), calculated as the relative mean fluorescence during the time duration following the peak stimulus-evoked response (120-400ms), decreased from the Control to the LED condition (Control vs LED p=0.016, Control vs Double LED p=0.020, paired t- test, n= 9 recordings). LED approx. 17mW/mm^2^ ramp, Double LED approx. 35mW/mm^2^ ramp – see Methods. Error bars represent mean +/- SEM.

To summarize, given the boosting of thalamic bursting induced by thalamic hyperpolarization that serves as the primary feedforward inputs to cortical S1, it was surprising to observe in the awake mouse not an increase, but instead a sensory evoked cortical response that was invariant in absolute amplitude and decreased relative to background activity.

### Thalamic Bursting and the Spatiotemporal Shaping of the Sensory Evoked Cortical Response

The GEVI imaging of cortex enables further investigation of the spatiotemporal characteristics of the observed phenomenon. Figure 6A shows the imaging frame associated with the peak sensory-evoked fluorescence, for the Control (top) and thalamic hyperpolarization (LED, bottom) conditions. What is apparent qualitatively from the images is that there was an overall attenuation of the cortical evoked response with thalamic hyperpolarization, as previously shown in Figure 4. Less immediately obvious is the effect of the thalamic hyperpolarization on the spatial extent of the sensory evoked cortical activation, as an overall loss in amplitude would trivially produce an apparent loss in spatial activation, as illustrated conceptually in the top panel of Figure 6B – often referred to as the “iceberg” effect. The bottom panel of Figure 6B shows conceptually the alternative - that the reduction in spatial area of activation may not just be due to the reduction in amplitude of the evoked response, but instead can reflect a spatial sharpening of the sensory evoked response with increasing thalamic hyperpolarization after accounting for the decrease in peak amplitude of fluorescence. For the GEVI recordings here, the latter is indeed the case - this is summarized as a reduction in the evoked normalized area of activation in the left panel of Figure 6D (Control vs LED p=0.025, Control vs Double LED p=0.046, paired t-test, 2 mice over 9 recording sessions for LED conditions).

In addition to the spatial characteristics of the sensory evoked response revealed by the voltage imaging, the voltage imaging is also useful in revealing relatively fast temporal dynamics of the aggregate cortical activity. Figure 6C shows the integrated fluorescence signal within a barrel column in the presence (LED condition) or absence (Control) of light input to activate halorhodopsin in thalamus, giving a picture of the temporal dynamics of the sensory-evoked clear response to the punctate whisker stimulus (2 mice over 9 recording sessions). We again found a decrease of the sensory evoked response (note that the fluorescence in Fig. 5C is normalized) despite a significant boosting of the sensory-evoked thalamic bursting (Fig. 4). Also apparent is the presence of a more rapid, post-peak decay in the fluorescence with thalamic hyperpolarization, and a prolonged period of sub-baseline fluorescence before returning to baseline after several hundred milliseconds. To quantify this property of the sensory evoked response, we calculated the temporal width of the evoked response as the time between the fluorescence peak and the first return to baseline level (defined as peak-to-baseline, Figure 6C), as shown in the middle panel of Figure 6D, revealing a clear narrowing/shortening of the temporal response as a function of thalamic hyperpolarization from ∼150ms in the Control condition, to ∼67ms in the thalamic hyperpolarized condition (Control vs LED p=0.0059, Control vs Double LED p=0.0019, unpaired t-test, LED conditions n=9, Control n=8 recording sessions, note one Control recording did not return to a pre-stimulus baseline). To quantify the prolonged post-stimulus undershoot of activity, we calculated the overall fluorescence in a window from 120 to 400ms post-stimulus (defined as mean recovery in Figure 6C, Control vs LED p=0.016, Control vs Double LED p=0.020, paired t-test, n=9 recording sessions), as shown in the right panel of Figure 6D. The increasingly negative relative fluorescence in this window revealed a clear increase in the undershoot with increasing thalamic hyperpolarization. Taken together, these S1 voltage imaging results show that in addition to the attenuation of the sensory-evoked response with thalamic hyperpolarization, there is a corresponding sharpening of the spatial activation of S1 and a temporal narrowing/shortening in the form of a more transient evoked response with an inhibitory undershoot.

### Thalamic Bursting and Cortical Spike Timing

Taken together, the results from Figure 6 revealed that the thalamic hyperpolarization strongly shapes both the spatial and temporal aspects of the cortical sensory evoked response in the awake brain. To further probe this phenomenon, we parsed our cortical single-units into putative excitatory neurons (regular spiking units, RSUs) and putative inhibitory neurons (fast spiking units, FSUs). Specifically, single-unit activity was identified (see Methods) and parsed based on RSU and FSU classification and combined across cortical layers, the result of which is shown in Figure 7A. Note that while this analysis involved combining recordings across the electrode array, we conducted a parallel analysis restricted to a smaller population of identified cortical layer 4 RSUs and FSUs (see below, and Supplemental Figure S5). The top panel of Figure 7 shows the waveform for typical putative RSU (red) and FSU (blue), where the solid line represents the mean waveform, and the band represents the standard deviation around the mean. The bottom panel shows the distribution of the time from the trough to the peak for the recorded waveforms. A threshold was set at 0.4 ms based on prior literature ((Guo et al., 2017), see Methods), for classifying the individual units into RSUs or FSUs.

**Figure 7.**
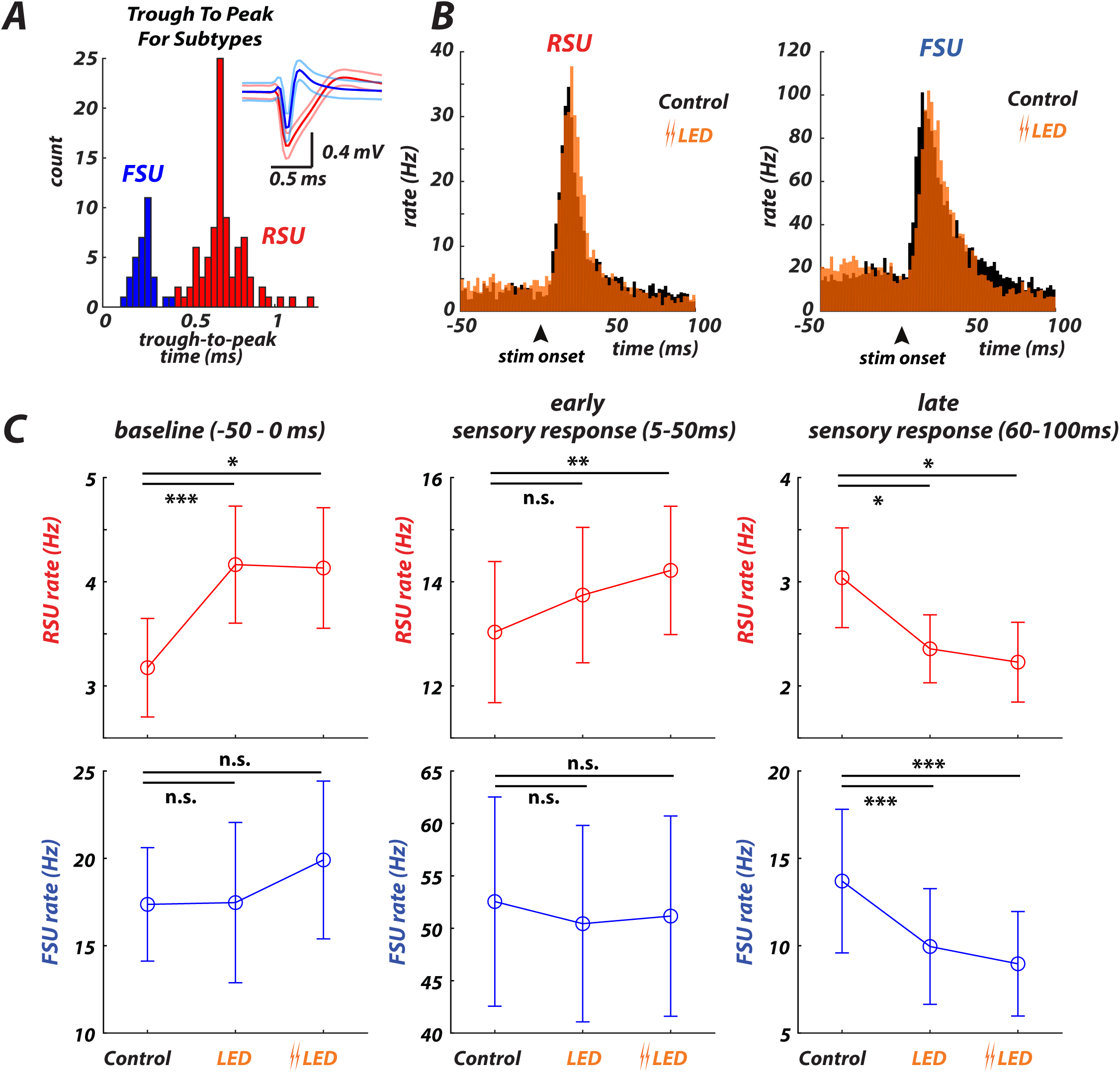
Thalamic hyperpolarization affects timing but not magnitude of sensory-evoked responses in cortex. **A.** Cortical single-units were classified as Regular Spiking Units (RSUs, red) or Fast Spiking Units (FSUs, blue) based on the time interval from the peak-to-trough (see Methods). Example RSU and FSU waveforms are shown at the top, where bands represent +/- 1 SD. The distribution of time intervals from peak-to-trough for the spike waveforms is shown at the bottom. **B.** PSTHs of the aggregate putative RSUs (left, n=86) and FSUs (right, n=32) in response to a punctate whisker deflection at time t=0 (bin size 2ms) for the Control (black) and double LED (orange) conditions. **C.** Summary analyses (Error bars represent mean +/- SEM) for RSUs (top row) and FSUs (bottom row). **Left** - Mean baseline (−50 to 0ms) RSU (Control vs LED p=5.3e-4, Control vs Double LED p=0.012, Wilcoxon signed-rank test, n=86) and FSU (Control vs LED p=0.15, Control vs Double LED p=0.30, Wilcoxon signed-rank test, n=32) firing rates for the LED relative to the Control condition. **Middle** - RSU (Control vs LED p=0.067, Control vs Double LED p=0.0082, Wilcoxon signed-rank test, n=86) and FSU (Control vs LED, p=0.30, Control vs Double LED p=0.45, Wilcoxon signed-rank test, n=32) early sensory response (5-50ms) for the LED compared to the Control conditions. **Right** – Late sensory response (60-100ms) for the LED compared to the Control condition for the RSU (Control vs LED p=0.017, Control vs Double LED p=0.014, n=86, Wilcoxon signed-rank test) and FSU (Control vs LED p=2.2e-4, Control vs Double LED p=4.5e-4, n=32, Wilcoxon signed-rank test) populations. LED approx. 17mW/mm^2^ ramp, Double LED approx. 35mW/mm^2^ ramp – see Methods. Error bars represent mean +/- SEM.

Apparent in the aggregate PSTH is the expected higher baseline/ongoing activity and sensory evoked responses in the FSUs compared to the RSUs, as shown in Figure 7B (∼17 HZ for FSU, n=32; ∼3 Hz for RSU, n=86), along with several other observations quantified in Figure 7C. As shown in the left panels of Figure 7C, in response to the thalamic hyperpolarization (LED ∼17 mW/mm^2^, double LED ∼35 mW/mm^2^), there was a relatively modest increase in baseline (pre-stimulus, -50 – 0 ms) firing rate for RSUs (Control vs LED p=5.3e-4, Control vs Double LED p=0.012, Wilcoxon signed-rank test, n=86), while FSUs showed no significant change (Control vs LED p=0.15, Control vs Double LED p=0.30, Wilcoxon signed-rank test, n=32). The response to the punctate sensory stimulus was a relatively short latency, transient increase in firing activity in both cell types, appearing to be relatively unchanged in overall magnitude with thalamic hyperpolarization, further quantified in the middle panels of Figures 6C (early response, 5-50ms post stimulus). We observed a slight increase in the RSU evoked firing rate (less than ∼10% increase, Control vs LED p=0.067, Control vs Double LED p=0.0082, Wilcoxon signed-rank test, n=86); however, FSUs displayed no statistical difference (Control vs LED p=0.30, Control vs Double LED p=0.45, Wilcoxon signed-rank test, n=32).

Although the overall magnitude of the early phase of the sensory evoked response was relatively unchanged by the thalamic hyperpolarization for both the FSUs and RSUs, there was a qualitative change in shape of the PSTH, with an increased latency to peak in the early phase, and change in the later phase of the evoked response. Specifically, thalamic hyperpolarization induced a reduction in the later phase of the cortical response, and a dip below the baseline firing rate in both the FSUs and RSUs, as shown in the PSTHs in Figure 7B. This was reflected in a decrease in the late phase (60-100ms) of the evoked response from the Control to the LED condition as shown in the right panels of Figure 7C (RSU - Control vs LED p=0.017, Control vs Double LED p=0.014, n=86, FSU Control vs LED p=2.2e-4, Control vs Double LED p=4.5e-4, n=32, Wilcoxon signed-rank test).

The preceding analysis was conducted by aggregating cortical units across probe recording sites, combining neurons across cortical layers. However, as cortical layer 4 is the primary recipient of synaptic input from VPm projections, we repeated the above analyses for putative cortical layer 4 neurons based on analysis of the multi-unit activity, local field potential, and corresponding current-source density (see Methods). In the cortical layer 4 analysis, we observe very similar results to those shown for the larger aggregate dataset, and the trends we observed were qualitatively the same as shown in Figure 7 (see Supplemental Figure S5).

The changes in timing and shape of the PSTHs of the cortical neurons suggest the possibility that thalamic hyperpolarization could affect synchronization within the cortical network. We analyzed synchrony across simultaneously recorded cortical single-units. Synchrony in firing across a pair of Neurons 1 and 2 is evaluated by examining the spike times of Neuron 2 relative to a particular spike of Neuron 1, across all spikes of Neuron 1, as illustrated in Figure 8A (see Methods). This forms the spike cross-correlogram (CCG), from which the synchrony is calculated as the integrated area within a +/- 7.5 ms window (see Methods, (Wang et al., 2010; Whitmire et al., 2016)), as illustrated in the bottom of Figure 8A. Due to the sensitivity of the synchrony metric to firing rate, we only examined pairs with a robust measurement (more than 50 synchronous events) to control for measurement accuracy (experimental results were invariant with different thresholds, see Methods), which restricted the analysis to FSUs in our dataset. Figure 8B shows rasters from an example pair of FSUs in the Control and double LED conditions (top), along with the spikes identified to be synchronous by this criterion (bottom). Figure 8C shows the aggregate cross-correlograms across 99 FSU pairs for the sensory evoked response, calculated from the spiking in the 100 ms window following the delivery of the sensory stimulus. There is an increase in the concentration of mass around 0 lag with thalamic hyperpolarization, which is indicative of an increased synchrony. This is summarized in Figure 8D, showing a significant increase across FSU units in the measured synchrony (Control vs LED, p=1.5e-8, Control vs Double LED, p=3.5e-13, Wilcoxon signed-rank test, n=99 FSU pairs). For comparison, the synchrony was also computed for the ongoing, spontaneous activity, revealing the synchronizing effect of the transient sensory input. These results were qualitatively similar for various synchrony window sizes (not shown).

**Figure 8.**
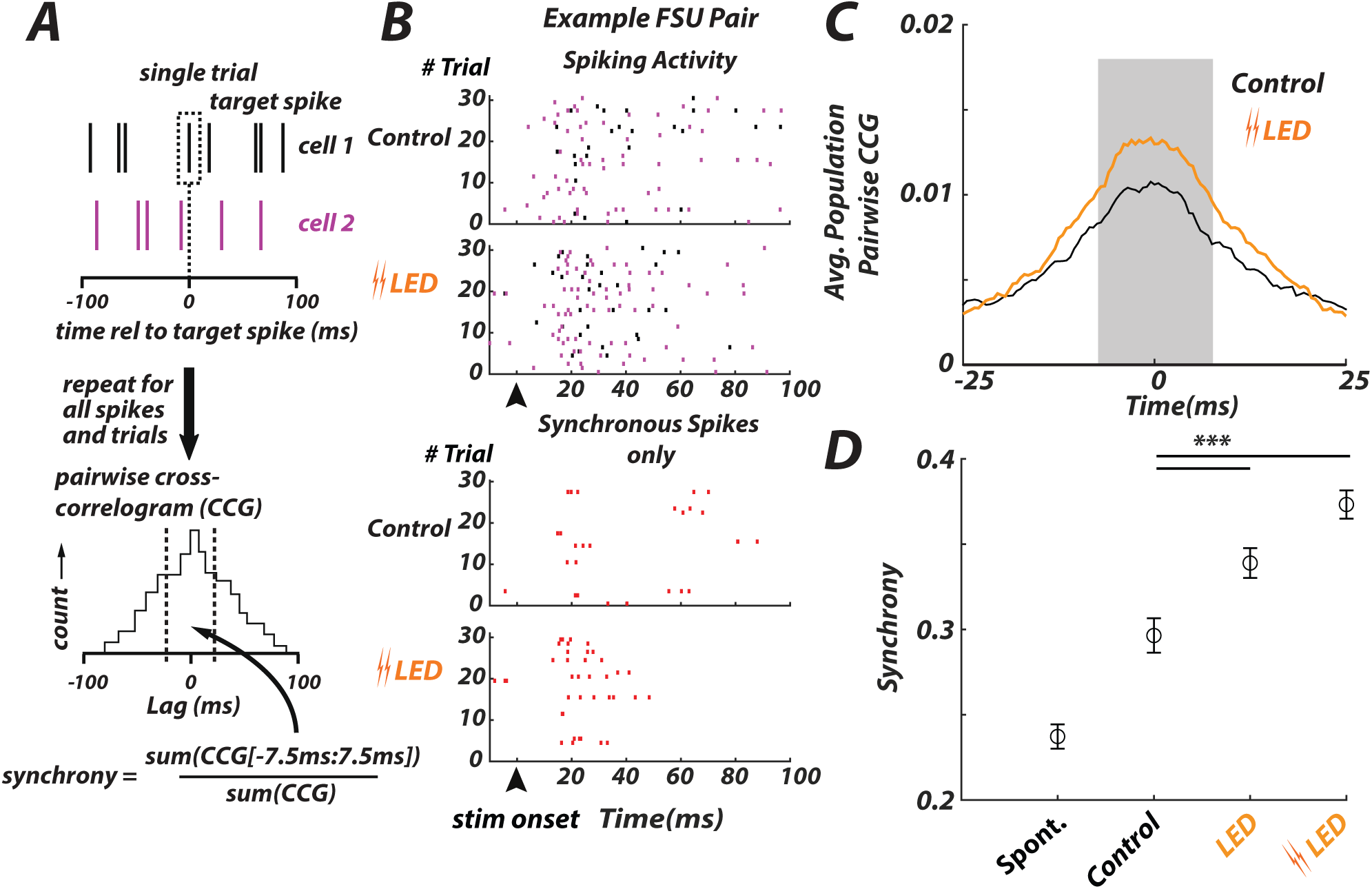
Thalamic hyperpolarization enhances cortical synchrony. **A.** The spike cross- correlogram (CCG) was calculated as a histogram of spike times of cell 2 relative to a target spike of cell 1, repeated for all spikes of cell 1. The pair-wise synchrony was calculated from the CCG as the area of the CCG between +/- 7.5ms, normalized by the total area of the CCG. **B**. **Top** Example Raster of two simultaneously collected FSU units, Cell 1 (black), Cell 2 (purple) for Control and LED conditions. **Bottom**. Synchronous spiking events only for the same neural pair (red, firing within 7.5ms of each other). **C.** Aggregate cross-correlograms for FSUs (n=99 pairs, see Methods) for the control (black) and LED (orange) conditions. Note that for the LED condition, the light level was 35 mW/mm^2^ to better emphasize the change in synchrony with thalamic hyperpolarization. Cross-correlograms have been smoothed via a moving average filter, 2.5ms window. **D.** Mean levels of synchrony FSU pairs, for the spontaneous/baseline cortical firing as compared to the sensory-evoked response for the Control and LED conditions. Relative to the Control condition, thalamic hyperpolarization (LED, 35mW/mm^2) resulted in an increase in synchrony for the FSU (Control vs LED, p=1.5e-8, Control vs Double LED, p=3.5e-13, Wilcoxon signed-rank test, n=99 pairs) populations. LED approx. 17mW/mm^2^ ramp, Double LED approx. 35mW/mm^2^ ramp – see Methods. Error bars represent mean +/- SEM.

### Modeling of the thalamic burst driven cortical E-I circuit

We next sought to understand the mechanistic basis of two key experimental results: 1) the nearly invariant absolute S1 sensory response in the thalamic hyperpolarized (LED) condition in the awake mouse, despite the increase in sensory-evoked burst spikes in VPm; and 2) the increase in sensory-evoked cortical synchrony in FS neurons in the thalamic hyperpolarized (LED) condition. To explore the potential role of various thalamic and cortical mechanisms, we constructed a simple model of the thalamocortical network, as described previously (Wright et al., 2021). This network mimics the numerical expansion of neurons at the thalamocortical junction, and incorporates several known properties of thalamocortical and intracortical connectivity (see Methods and Supplemental Information on Cortical E-I Modeling). We previously tuned model parameters to produce qualitatively realistic velocity tuning curves (Wright et al., 2021), using those parameters here. The network was composed of a single model VPm “barreloid” (40 independent spike trains) projecting to a single cortical barrel “column”, modeled as an interconnected network of 800 excitatory and 100 inhibitory single-compartment leaky integrate-and-fire neurons with clustered connectivity (Litwin-Kumar and Doiron, 2012; Bujan et al., 2015; Wright et al., 2017b, 2017a). To create the model input, ongoing and sensory-evoked VPm tonic and burst spikes were generated based on the empirical VPm PSTHs (Figure 9A, top two rows, see Methods). Generating thalamic input spikes in this manner qualitatively reproduced the high rate of synchronous spikes in the short-latency response window. Across conditions the rate of synchronous spikes in the model VPm input matched those of our experimental observations, confirmed using the approach described above for the empirical VPm data, with a slight but significant decrease in the synchronous spiking in the short-latency response window from Control to LED (Fig. 9B), consistent with what we observed experimentally. Because we sought to investigate the effects of such changes in synchronous thalamic spiking and single-neuron bursting on downstream network activity, we did not impose activity-dependent depression at the thalamocortical synapse. Given the very short intervals between burst spikes, the relatively slow recovery of a given thalamocortical synapse following the first spike in a burst would in principle diminish the efficacy with which subsequent spikes in the burst drive postsynaptic targets. Yet we show below that this consideration is not required to match experimental observations. For a more detailed description of the model, see Supplemental Information on Cortical E-I Modeling.

**Figure 9.**
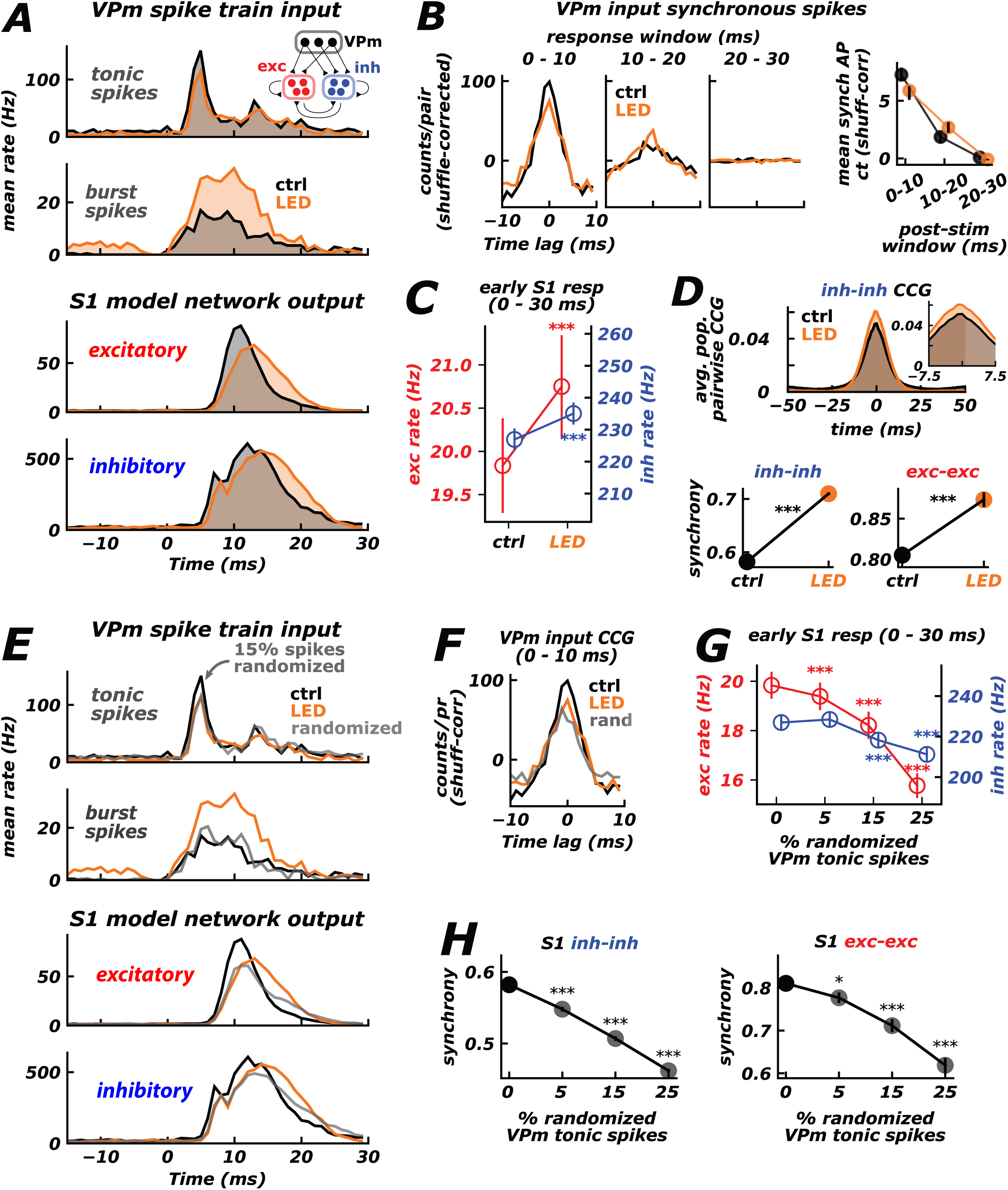
A thalamocortical network model reproduces key experiment results implicating thalamic spike timing. **A.** Thalamocortical model schematic (see Methods). **Top inset** - A clustered network of excitatory (red) and inhibitory (blue) neurons received inputs from a model VPm barreloid, as well as random excitatory inputs. **Top** - Grand mean PSTHs for VPm tonic and burst spikes. **Bottom** – Grand mean PSTHs for network excitatory and inhibitory neurons, for the Control (black) and LED (orange) conditions. **B. Left** – Grand spike cross- correlograms for VPm spike train inputs to the model cortical network, for three post-stimulus windows. **Right** – Mean synchronous spike counts for VPm spike train inputs to the model cortical network. Error bars indicate 95% confidence intervals from re-sampling relative spike times with replacement. **C.** Grand mean (+/- SEM) rates for all model network excitatory (red) and inhibitory (blue) neurons for the early (0 – 30 ms post-stim) response window. Excitatory control vs LED: 4.6% increase, p = 6.85 x 10^-9^, Wilcoxon signed-rank test. Inhibitory control vs. LED: 3.6% increase, p = 2.15 x 10^-13^, Wilcoxon signed-rank test. **D.** Top – Grand cross- correlograms for 200 randomly-selected, valid pairs of network inhibitory neurons (see Methods). **Inset** – Same, but for window used to calculate synchrony (+/- 7.5 ms). **Bottom** - Grand mean (+/- SEM) pairwise synchrony for 200 valid inhibitory pairs (left) and 147 valid excitatory pairs (right). Inh-inh control vs. LED: 22.4% increase, p = 1.44 x 10^-34^, Wilcoxon signed- rank test. Exc-exc control vs. LED: 8.4% increase, p = 1.74 x 10^-7^, Wilcoxon signed-rank test. **E**. Same as in (A), but including “randomized” condition (gray PSTHs), in which P = 15% of drawn tonic spike times occurring near the early PSTH peak were shifted to a later, random time. **F.** Same as in (B, left), but including “randomized” condition (gray CCG). **G.** Same as in (C), but for various choices of P (percent randomized VPm tonic spike times). **H.** Same as in (D, bottom), but for various choices of P. * Indicates 0.01 ≤ p < 0.05, ** indicates 0.001 ≤ p < 0.01, *** indicates p < 0.001, Wilcoxon signed-rank test.

This model network succeeded in qualitatively reproducing the two results identified above. First, despite the substantial increase in burst spikes in the thalamic input driving the model cortical network (constructed to match empirical rates – Figure 9A, top), the network cortical response was relatively invariant with thalamic hyperpolarization, with only a very slight increase for the LED condition (Figure 9A, bottom two rows, and Figure 9C). Note that unlike the experimental data, synchrony measures from simulated cortical activity were not limited by lack of firing events, enabling synchrony measurements across both excitatory and inhibitory sub-populations. We found that the evoked pairwise synchrony of S1 neurons was higher in the LED condition for both excitatory and inhibitory model neurons (Fig. 9D). In short, the mechanisms incorporated in this simple model were sufficient to predict the two key experimental results.

Having reproduced these results, we next sought to test the hypothesis that the nearly invariant mean cortical response despite the boosting of thalamic bursting represented the net effect of two opposing mechanisms: enhanced thalamic bursting in the LED condition supporting more robust thalamocortical synaptic drive, counterbalanced by a reduction in the rate of spikes that are synchronous across thalamic neurons. To this end, we manipulated the rate of synchronous thalamic input spikes, while maintaining the mean rate of tonic and burst spikes, analogous to our previous work (Wang et al., 2010; Wright et al., 2021). Specifically, an artificial condition was simulated in which tonic spike times were probabilistically shifted away from the early PSTH peak, such that the peak firing rate of this manipulated condition approximately matched that of the LED condition, referring to this manipulation as “randomized” (Fig. 9E, top). The grand VPm input CCG was again generated, confirming that this manipulation reduced the rate of synchronous spikes in the input spike trains (Fig. 9F). Because the rate of tonic spikes in the 30 ms post-stimulus window was not altered and used the burst spike distribution from the Control condition, this enabled the effects of such changes in synchronous spiking alone to be inferred, without a possibly counterbalancing increase in bursting. In contrast to the LED condition, shifting even small percentages of tonic VPm spikes (i.e., reducing the rate of synchronous VPm spikes) resulted in a significant decrease in cortical network response rates (Fig. 9G) and synchrony (Fig. 9H), with both measures decreasing monotonically with increasing percentage of shifted VPm spikes. These simulations thus support the notion that while thalamic bursting per se likely provides highly efficacious synaptic drive to cortex, the loss of short-latency, synchronous tonic firing in the LED condition counterbalances this effect, resulting in a nearly invariant S1 response. Further, the trends observed in cortical synchrony (Fig. 9H) suggest thalamic bursts play an important role in synchronizing cortical neurons.

Taken together, the modeling results support the notion that cortical sensory representations are highly sensitive to thalamic firing. The degree of i) thalamic bursting in individual neurons and ii) synchronous spiking across the thalamic population together determine the efficacy with which VPm spiking drives S1. This predicts the counterintuitive empirical result that the enhanced VPm bursting induced by hyperpolarization does not result in a substantially boosted S1 response.

## DISCUSSION

The sensory thalamus controls the flow of signaling from the periphery to cortex, ultimately gating what we do and do not perceive about the outside world. Despite its critical role in sensing, how this circuit controls signaling remains poorly understood. Here, through a range of experimental approaches in the awake, head-fixed mouse, we show that optogenetic thalamic hyperpolarization significantly enhances sensory-evoked bursting, yet the baseline thalamic firing rate and sensory evoked magnitude are both relatively invariant. Sensory cortex subsequently exhibits a surprisingly invariant absolute evoked response despite the potent thalamic burst input, instead demonstrating increased timing precision, a focusing of spatial activation, and increased synchrony of spiking. Thalamocortical network modeling further supports the assertion that the bursting-induced changes in thalamic spike timing and thalamic population synchrony are sufficient to explain the increase in cortical synchronization and the invariant cortical response amplitude, respectively. The findings here present a highly sensitive, timing-based gating of sensory signaling to cortex.

### Thalamocortical Gating of Sensory Signals

One surprising observation was that the absolute cortical sensory evoked response amplitude was invariant to thalamic hyperpolarization in awake mice, despite a significant increase in thalamic bursting. At first glance, this appears to contradict previous work clearly demonstrating the increased efficacy with which bursts in individual thalamic neurons impact monosynaptically-connected cortical neurons at the single synapse level (Swadlow and Gusev, 2001). Importantly, however, the efficacy with which a population of thalamic neurons drives cortical spiking depends on relative spike timing both within a given thalamic neuron (i.e., tonic vs. burst) and across convergent thalamic neurons (i.e., degree of synchronization). We found that while thalamic hyperpolarization promoted single-neuron bursting, there was a surprising decrease in synchronous thalamic firing in the critical early sensory-evoked response. The nearly invariant cortical response makes sense in light of this observation; cortex is extremely sensitive to thalamic spike timing over this 10ms period established by the disynaptic inhibition mediated “window of opportunity” (Pinto et al., 2000; Wilent and Contreras, 2004, 2005; Gabernet et al., 2005). The findings here thus coexist with the potency of thalamic bursts in driving post-synaptic cortical responses at the single synapse level.

While the absolute cortical response amplitude was nearly invariant to thalamic hyperpolarization, the timing precision and synchrony of the cortical response were enhanced by our manipulation. Such changes in cortical representations suggest a more potent input for subsequent downstream signaling, and perhaps enhanced stimulus detectability - despite the invariance of S1 firing rate – but corresponding recordings of the S1 recipient regions and a behavioral assay would be needed to test this hypothesis directly. We also note that the observed changes in FS synchrony likely play a role in shaping the overall cortical response we observe, as well as shaping the cortical sensitivity to timing of the thalamic input through direct interactions with the excitatory cortical sub-population, but a full exploration of the excitatory/inhibitory interactions within cortex was beyond the scope of what was investigated here. Further, the nature of the short ISIs within thalamic bursts almost certainly means that depression at the thalamocortical synapse plays a role in determining the potency of the sensory-evoked cortical response, as has been previously shown. However, the modeling results here suggest that single-neuron bursting and across-neuron synchronous thalamic firing, and how they interact with the timing sensitivity of cortex established by the disynaptic feedforward inhibition, are the key role players in the observed dynamics, although more extensive experiments involving causal manipulations of these mechanisms would need to be conducted to more conclusively establish this explanation. Finally, though not explicitly tested in these experiments, the results here would suggest that important coding aspects of this particular pathway, notably velocity and direction tuning, would be strongly shaped through changes in the timing properties of the thalamic population, as we have previously shown in the context of rapid sensory adaptation (Wang et al., 2010).

In several of the analyses here, we combined recorded cortical units across layers. While the optical GEVI imaging approach is focused on S1 layer 2/3, it also likely captures activity above and below layer 2/3, as well as any cross-laminar processes, and further obscures any differences across excitatory and inhibitory sub-populations due to the indiscriminate viral targeting. Neurons in VPm thalamus send axonal projections differentially across laminae in S1 (Sermet et al., 2019), likely resulting in variations in the effects of thalamic hyperpolarization on S1 neurons in different cortical layers. To account for the possibility that the pooled dataset was not representative of thalamorecipient neurons, we repeated a subset of our analyses for putative S1 layer 4 neurons, and in each case observed the same trends as in the aggregate data (Supplemental Figure S5). Note that the identification of L4 neurons was conducted conservatively, and thus some of the non-L4 neurons in the aggregate population are likely also L4. Importantly, the putative L4 neurons had a larger sensory evoked response than the non-L4 neurons, as is apparent in comparing Supplemental Figure S5 to Figures 5 and 7, thus likely dominating the aggregate analysis. Further, note that while the measurements of synchrony in S1 here were limited to the putative FS neurons due to the relatively low firing rates of the RS neurons, the modeling results suggest that the RS neurons would also exhibit an increase in synchrony with thalamic hyperpolarization, but this must be directly measured in future studies. Regardless, the qualitative consistency of our cortical results across cell types, laminar location, and recording modalities suggests that our observations provide a representative – though by no means comprehensive – view of the net effects of thalamic spike timing on cortical sensory responses.

### Invariance in Baseline Thalamic Firing Activity

In this study, we utilized optogenetic hyperpolarization to bias thalamic sensory responses toward bursting, without significantly changing other dynamics that might indirectly impact sensory responses. One particularly surprising finding here was that despite optogenetically induced hyperpolarization of thalamus, the overall baseline firing rate of thalamic neurons was not suppressed, but instead unchanged. Following a transient decrease in firing rate, the recorded VPm neurons returned to their original baseline firing rate. This finding is, however, consistent with other reported observations. In the visual pathway for example, optogenetic excitation of the thalamic reticular nucleus (TRN) transiently silences the lateral geniculate nucleus, followed by a return to the original firing rate at steady-state (Reinhold et al., 2015). In the somatosensory pathway, strong photoinhibition attenuated VPm thalamic firing rate, but failed to quench activity altogether (Halassa et al., 2011; Poulet et al., 2012; Lewis et al., 2015; Reinhold et al., 2015; Yu et al., 2016). The most parsimonious explanation given the observations here is that moderate amounts of hyperpolarization serve to engage the dynamics of the T-type calcium channels that are inactive at normal baseline conditions, effectively compensating for the loss in tonic spiking due to the hyperpolarization. Although the optogenetic approach here does not enable direct observation of the magnitude of the hyperpolarizing input, separate intracellular in-vitro slice experiments where we repeated the protocol while patching on to VPm neurons revealed relatively modest amounts of hyperpolarization that were well-sustained during constant light illumination. This is important, as recent studies have suggested that prolonged activation of specific opsins can have unintended consequences, notably here the possibility of changes in the reversal potential for chloride (Raimondo et al., 2012). This effect could theoretically result in changes in the degree of hyperpolarization, although halorhodopsin as a pump is less directly affected by immediate (local) changes in reversal potential as compared to channel-based optogenetics. Further, with halorhodopsin there is the potential for photoinactivation and decreased photocurrents (Zhang et al., 2019), which could also change the degree of hyperpolarization over time. Through the combination of the intracellular control experiments and replication of the primary result through the integrate and fire or burst (IFB) model, the likelihood that these possible effects played a primary role here is low, especially over the relatively short timescales considered here.

Importantly, beyond the engagement of the T-type calcium channel burst mechanism upon initial hyperpolarization, it would seem that additional hyperpolarization would further push the neuron away from threshold, making it more difficult to burst, which would predict a corresponding decrease in firing rate. However, we found that increased hyperpolarization increased the number of spikes per burst, which served to offset the decrease in the number of bursts with increasing hyperpolarization. The result is a surprisingly resilient mechanism in response to this perturbation. It should be noted that other elements of the circuit likely play a role in the observation here – for example, the initial decrease in VPm firing rate would decrease excitation of TRN, subsequently decreasing inhibition of VPm, which would work synergistically with the intrinsic properties of the burst mechanism in this compensatory action.

### The Potential Role of Thalamic Bursts in Sensory Signaling

Almost four decades ago, Crick proposed a provocative hypothesis – that the thalamic- reticular complex serves as a dynamic gate for attentional control of sensory signaling to cortex (Crick, 1984). Further refinement of this idea suggested that the switching between tonic and burst firing modes of thalamic neurons that is facilitated by the unique dynamics of the T-type calcium channels that are prevalent in the thalamus could establish a context dependent signaling (Sherman and Guillery, 2002; Sherman, 2005). In this framework, burst spiking would promote the detection of salient sensory features, while tonic spiking would promote the transmission of details about the nature of the sensory stimulus. Furthermore, the thalamic burst would also potentially provide a “wake-up call” to cortex (Sherman, 2001), garnering attentional resources that ultimately would serve to switch the thalamic mode of firing to tonic through depolarizing corticothalamic feedback. As attractive a framework as this is, it is also a daunting theory to test experimentally due to the complexity of the circuit, the required specificity of recording and manipulation, and ultimately the need to cast in the context of changing states of arousal during trained behaviors. Previous anesthetized studies have made substantial progress toward this goal. For example, sensory-evoked thalamic bursting under anesthesia has been shown to be well-driven by the appearance of salient sensory features (Lesica and Stanley, 2004; Alitto et al., 2005; Denning and Reinagel, 2005), which promotes the detection of change in the sensory input from the perspective of an ideal observer of thalamic spiking (Lesica and Stanley, 2004), and this sensitivity is strongly shaped by thalamic state (Lesica et al., 2006). Further studies showing the sensitivity of sensory cortex to spontaneous (non-sensory) thalamic bursting during wakefulness (Swadlow and Gusev, 2001) seem to set the stage for at least part of the overarching coding scheme in which the cortical response would be amplified by the thalamic bursts in “detect” mode.

In opposition to this view, some have noted that because ongoing thalamic spontaneous bursting events are particularly prominent during slow wave sleep and under anesthesia, they likely play no role in sensory signaling during wakefulness. However, this has been largely disproven, with low rates of bursts occurring both spontaneously and during naturalistic stimuli in awake somatosensory (Swadlow and Gusev, 2001; Stoelzel et al., 2009; Wright et al., 2021), visual (Niell and Stryker, 2010), and auditory systems (Massaux et al., 2004). Further, we have shown previously in the awake, head-fixed rat (Whitmire et al. 2016) and mouse (Wright et al. 2021) that rapid sensory adaptation has a particularly strong effect on sensory-evoked burst firing, suggesting that the adaptive changes in bursting may play an important role in perceptual adaptation during wakefulness. Thus, it is critical to precisely probe the effects of sensory-evoked thalamic bursting on cortical sensory representations during wakefulness, and specifically to test the decades-old “wake-up call” hypothesis.

To this end, an optogenetic manipulation approach was adopted to gain systematic control over thalamic burst/tonic firing modes, and the functional effects were precisely quantified at the level of thalamus and primary sensory cortex. During normal physiological conditions, the thalamus receives a range of complex, excitatory and inhibitory inputs that interact with the intrinsic cell properties to collectively set the baseline membrane potential and firing rate of these neurons as a function of behavioral state. Cortex also exhibits profound changes in ongoing and sensory-evoked firing across states of wakefulness, and there is strong evidence that thalamic activity itself is a strong driver of cortical state (Poulet et al., 2012). But because thalamus and cortex are densely interconnected through feedforward and feedback projections, and are both subject to a range of other modulatory inputs, it is generally difficult to infer directions of causality. The approach here enabled the disentangling of these interactions by probing the downstream effects of sensory-evoked thalamic bursting and synchronous firing per se, independent of the many other naturally-occurring fluctuations. We do not suggest that the thalamic hyperpolarization mimics a particular behavioral state, but rather that it provides an opportunity to bias thalamic sensory responses toward burst firing mode, so that we might probe the consequences for cortex. Such targeted manipulations are critical to understanding state-dependent sensory signaling in this complex circuit more generally.

In the experiments presented here, in the absence of optogenetic manipulation, bursting represents approximately 15% of the total sensory evoked response, as compared to greater than 30% during imposed thalamic hyperpolarization. Thus, even in the absence of optogenetic manipulation, the bursting is not insignificant, and determining the effects of thalamic bursts on cortex is relevant even under normal conditions. It is also important to note that awake, head-fixed mice likely explore a much narrower range of behavioral (e.g., arousal) states than occur in naturalistic settings. It is well-known that states of low-arousal are directly correlated with thalamic burst firing modes, thus predictive of more sensory-evoked burst firing than we observed here. While the experimental preparation used here provides the requisite stability to perform precise stimulation and recording, a complementary scenario for exploring the role of bursting across states of wakefulness would involve freely moving rodents across a wider range of behavioral states. These kinds of studies, in concert with those providing even more precise control of thalamic firing modes (e.g., via closed-loop feedback control of neural activity (Bolus et al., 2018, 2021)) need to be employed to more strongly, effectively, and comprehensively explore the coupling across brain regions and behavioral states.

Although the results here in the awake brain demonstrate that the absolute S1 sensory evoked response was invariant to thalamic hyperpolarization and even diminished relative to background cortical activity despite potent sensory-evoked thalamic bursting, what emerges is increased timing precision, increased focus of spatial activation, and a corresponding synchronization of the cortical sensory-evoked response that collectively could promote detectability. Given the likely timing sensitivity of down-stream brain structures in the sensorimotor arc, the synchronization of cortical activity may in fact be a more critical element of cortical signaling than overall magnitude, supported by behavioral work demonstrating the importance of cortical synchrony over firing rate (Jadhav et al., 2009). Taken together, the results here point to timing rather than response magnitude as a fundamental currency of the thalamocortical circuit, presenting a dynamic, timing-based gating of sensory signaling to cortex that has strong implications for detectability and discriminability in complex sensory environments.

## Acknowledgements

This work was supported by NIH National Institute of Neurological Disorders and Stroke Grant R01-NS048285 (GBS), R01-NS104928 (GBS), U01-NS094302 (GBS and DJ), and NIH National Institute of Neurological Disorders and Stroke Pre-doctoral NRSA NS098691 (PYB). The authors would like to thank Katie Borden, Ilya Kolb, Will Stoy, and Precision Biosystems Lab at Georgia Tech, for additional support and advice on analysis. Additionally, we thank all members of the Stanley Laboratory for the Control of Neural Systems at Georgia Tech for feedback and experimental design which made this work possible

## STAR METHODS

### Resource Availability

#### Lead Contact

Further information and requests for data or code should be directed to and will be fulfilled by the lead contact, Garrett Stanley.

#### Materials Availability

This study did not generate new unique reagents.

#### Data and Code Availability

All data reported in this paper will be shared by the lead contact upon request.

All original code will be deposited in a public repository and will be publicly available as of the date of publication. DOIs will be listed in the key resources table.

Any additional information required to reanalyze the data reported in this paper is available from the lead contact upon request.

### Experimental Model and Subject Details

All procedures were approved by the Institutional Animal Care and Use Committee at the Georgia Institute of Technology and were in agreement with guidelines established by the National Institutes of Health. Experiments were performed on adult (6 – 15 weeks) male and female Mice. Mice strains included C57BL/6J (Jackson Laboratories) and the transgenic NR133-Cre (Gerfen et al., 2013) on a C57BL/6J background mice.

### Method Details

#### AAV Delivery

At least 5 weeks prior to experimentation, 6 week old female C57BL/6J (Jackson Laboratories) mice were injected with different viral constructs either in the ventral posteromedial (VPm) thalamic region with AAV-5-CamKinaseII-eNph3.0 (UNC Viral Vector core) for optogenetic modulation, in the primary somatosensory (S1) cortex with AAV-1-hsyn1-ArcLight (UPenn Viral Vector Core) for optical voltage imaging, or both. For surgical procedures, mice were anesthetized using isoflurane (3-5%). After the mouse was fully anesthetized, small craniotomies were placed over the regions of interest and were aligned using stereotaxic measurements (For VPm, 1.8mm lateral from midline by 1.8mm caudal from bregma). For cortical expression, either single or multiple injection sites were used surrounding the barrel cortex (centered at 1.5mm caudal from bregma and 3mm lateral from midline). The virus was loaded into a modified Hamilton syringe (701-N) with a ∼35 micron borosilicate glass pipette tip or a Hamilton Neuros Syringe. The syringe was initially lowered to the corresponding depth below the surface (for VPm: 3mm and For S1: 0.5mm) and the tissue was allowed to rest for 1 minute before injection. Both sites received injections of 0.5-1µl of viral construct at a flow rate of 0.1µl/minute. After injection, the pipette remained in place for an additional 5 minutes before slowly being removed from the brain. The bore holes were filled with either bone wax or left to close naturally. Throughout injection, mice were kept warm using a water heating system to maintain body temperature. See Supplemental Figure S1 for histological validation of expression of halorhodopsin in thalamus and ArcLight in S1.

#### Awake Animal Preparation

At least four weeks after ArcLight and eNphR3.0 viral injection, mice were anesthetized under isoflurane and were implanted with a head-plate. Over the course of 5-14 days preceding the first imaging experiment, mice were routinely handled to gain familiarity with the imaging system and immobilization device. During this acclimatization period, mice were head fixed for increasingly longer periods of time, from 15 minutes to 1.5 hours. During stimulation of the whisker, mice were prevented from interacting with the whisker stimulator by obstructing the path from the paws to the whisker. Mice were rewarded with sweetened milk (Nestle, Ltd.) throughout imaging. After at least 5 days of handling and acclimating, mice became tolerant to immobilization in the headplate restraint system. During passive stimulation of the whiskers, the mice often actively moved their whiskers. Therefore, the whisker stimulator was placed relatively close (5mm) to the face to prevent the whisker from slipping out of the manipulator; however, the amplitude of the deflection was adjusted to maintain a consistent angular velocity (1200 deg/s).

#### Whisker Stimulation

Whisker stimulation was similar to that utilized previously (Borden et al., 2017). Briefly, individual vibrissae of the mice were deflected by a high fidelity (1 KHz) galvanometer system (Cambridge Technologies). A whisker stimulus was applied by positioning the custom designed galvanometer 5-10mm from the face and delivering an exponential sawtooth (rise and fall time = 5ms). The waveform stimulus velocity was taken by averaging the time to peak velocity of the stimulus. The velocity was adjusted based on distance from the face.

#### Thalamic and Cortical Electrophysiology

For thalamic electrophysiology, a small craniotomy was made over the primary whisker sensitive thalamic ventral-posterior medial (VPm) region of the mouse, around the injection site, using stereotactic coordinates (see above). VPm was then mapped under anesthesia using either a 2MOhm tungsten electrode (FHC) or 32-channel silicon probe (NeuroNexus Technologies). The mapping electrode was slowly lowered below the cortical surface during manual stimulation of the whiskers, while spiking activity (threshold crossings of high-pass filtered voltages) was monitored using the data acquisition system. We stopped descending when we recorded whisker-driven spiking activity at a depth consistent with VPm (typically 2900 – 3600 µm below the surface). We then used the galvanometer to present precise single-whisker stimulation, and confirmed that multi-unit activity was consistent with electrophysiological features of VPm. Specifically, the electrode was determined to be located in VPm if the peri-stimulus time histogram (PSTH) contained a peak response 3ms - 10ms after a 1200 degree/s punctate single whisker stimulus and did not have a latency shift by more than 20ms after 1s of a 10Hz adapting stimulus (Wang et al., 2010). The principal whisker was determined by the largest 30ms PSTH response of multiple neighboring whiskers. We then noted the depth and stereotactic coordinates of this recording site and slowly retracted the mapping electrode, sealed the craniotomy, and returning the mouse to its home cage. During awake recording sessions, we targeted the same location for recording, briefly repeating the above steps to confirm electrode location. We recorded thalamic spiking using either a 2MOhm tungsten electrode (FHC) with 200 µm attached optical fiber, or a 32-channel silicon probe array with 100 µm attached optical fiber (A1x32-Poly3-5mm-25s-177- OA32LP, NeuroNexus Technologies). For cortical recordings, initial mapping was conducted using cortical ArcLight voltage imaging or intrinsic imaging (see below). Once the target cortical column (barrel) was identified and confirmed, a 32-channel linear silicon probe (A1x32-5mm-25-177, NeuroNexus Technologies) or single tungsten electrode (FHC) was inserted. For both thalamic and cortical electrophysiology, after the conclusion of the study either a small 7uA 10s lesion, or a fluorescent dye was placed near the recording location and confirmed using post-mortem histological validation. Neuronal signals were band-pass filtered (500Hz–5KHz), digitized at either 24.414 or 30 KHz/channel and collected using either a 96-channel (Blackrock Microsystems, Salt Lake City, UT, USA), or a 64-channel (Tucker-Davis Technologies, Alachua, FL, USA) data-acquisition system.

#### Awake Cortical Fluorescent ArcLight Imaging

ArcLight transfected mice were imaged through the thinned or removed skull using a two camera system: a Scimedia Imaging system to measure cortical ArcLight spatial activity, and a custom camera to measure hemodynamic activity for subtraction. The cortex was imaged using a 184 x123 pixel CCD Camera, MiCam2 HR Camera (Scimedia, Ltd) to capture ArcLight, and a Basler Ace (acA1920-155um) 480 x 180 pixel (4×4 binned) CMOS Camera to capture auto-fluorescence, at 200 Hz with a tandem lens microscope. The entire cortical area was illuminated at 465 nm with a 400 mW/cm2 LED system (Scimedia, Ltd.) to excite the ArcLight fluorophore and background auto-fluorescence. The excitation light was projected onto the cortical surface using the first dichroic mirror (bandpass: 475/625nm, Semrock, Inc.). Collected light was passed through a second dichroic mirror (Longpass cutoff: 495 nm, Semrock, Inc.) for collection of the ArcLight and auto-fluorescence signal. The auto-fluorescence signal was filtered with a bandpass filter between the wavelengths of 465/75 nm (Semrock, Inc). The ArcLight signal was filtered with a bandpass emission filter between wavelengths of 520⁄35 nm (Semrock, Inc.). The imaging system was focused approximately 300µm below the surface of the brain to target cortical layer 2/3, although the imaging likely captures fluorescence from the cell bodies as well as neuropil above and below layer 2/3.

#### Anesthetized Cortical Fluorescent ArcLight & Intrinsic Imaging

ArcLight transfected mice were imaged through the thinned or removed skull using a Scimedia Imaging system to measure cortical spatiotemporal activity (leveraging a single camera setup). The cortex was imaged using a 184×123 pixel CCD Camera, MiCam2 HR Camera (Scimedia, Ltd) at 200 Hz, and a tandem lens macroscope. The entire cortical area was illuminated at 465 nm with a 400 mW/cm^2^ LED system (Scimedia, Ltd.) to excite the ArcLight fluorophore. The excitation light was further filtered (cutoff: 472-430 nm bandpass filter, Semrock, Inc.) and projected onto the cortical surface using a dichroic mirror (cutoff: 495 nm, Semrock, Inc.). Collected light was filtered with a bandpass emission filter between wavelengths of 520-535 nm (Semrock, Inc.). The imaging system was focused approximately 300µm below the surface of the brain to target cortical layer 2/3. For intrinsic imaging of the hemodynamic response, the cortical surface was illuminated by a 625nm red LED (ThorLabs), and imaged with the same camera system as above, at a temporal resolution of 10Hz. During intrinsic imaging, no emission filters were used. In order to evoke a cortical intrinsic response, the whisker was repetitively stimulated at 10Hz for 6 seconds.

#### Functional Fluorescent Mapping of Barrel Cortex

The mouse’s whisker system was first mapped by imaging the rapid ArcLight response to a high velocity (1200 Deg/s) sensory stimulus separately applied to three different whiskers. The resulting whisker response averaged over 20 trials was determined to be associated with a principal whisker, and barrel, if the evoked response was spatially limited to roughly a 0.2 mm x 0.2 mm area 25-30ms after stimulation. Additionally, the response was determined to be originating from the barrel field if the center of mass of activation moved consistently with the histologically defined barrel field and was within the standard stereotaxic location of S1 (∼3mm lateral, 0.5-1.5mm from bregma). After mapping, a single whisker was deflected in a way as to emulate a high velocity slip-stick event (1200 deg/s), either with or without thalamic optogenetic hyperpolarization.

#### Simultaneous Imaging and Thalamic Optogenetic Manipulation

After mapping both the thalamic and cortical regions, an optrode (2M Ohm tungsten electrode mounted to a 200 µm optic fiber) was positioned to the stereotaxic locations of the pre-mapped thalamic region and lowered to the corresponding depth. Once a single thalamic unit was identified using the above constraints, the unit was determined to be sensitive to optical stimulation by briefly (1-2s) hyperpolarizing the cells using ∼17mW/mm^2^ (LED condition) or ∼35mW/mm^2^ (double LED condition) (unless otherwise noted) at 590nm from an LED light source (Thorlabs, M590-F1). Each cell was determined to be a thalamic optically sensitive unit if the lighted caused a transient decrease in firing rate or if the cessation of the 590nm light caused a rebound burst (Brecht and Sakmann, 2002). After identifying an optically sensitive thalamic unit, the whisker stimulus was presented under various light conditions. Light stimulation was presented 500-750ms preceding and following whisker deflection. There was at least a four second interval between stimulus deliveries to allow for recovery of halorhodopsin (eNphR3.0). Each session imaged 200ms-1s of frames preceding whisker stimulation to measure spontaneous activity. Prior to use, light power was measured from the tip of the ground optical fiber before each experiment to maintain approximate light intensities delivered to each cell. During light delivery, the downstream cortical response was recorded using either electrodes for cortical electrophysiology or voltage imaging as described above. The optogenetic and viral expression of each experiment was verified through confocal and brightfield imaging of fixed slices.

The LED light intensity used for optogenetic stimulation is within the published range of light stimulation (35mW/mm^2^ is estimated from our maximum power measure of ∼1.1mW through a 200µm fiber) (Stujenske et al., 2015; Owen et al., 2019). Further, studies that have directly quantified the effects of optical stimulation on local tissue heating and neural activation (in the absence of opsin expression) have found no significant difference in the firing rate change for 1mW light intensity (Stujenske et al., 2015) or minor firing rate changes for 3mW light intensity, but no behavioral effects (Owen et al., 2019). See Supplemental Figure S4 for controls that demonstrate a lack of light effects and lack of confounding interactions between optogenetic activation of VPm thalamus and GEVI imaging in cortical S1.

#### Anesthetized Electrophysiology

A subset of experiments was conducted with mice under light anesthesia, as a control. These mice were initially anesthetized using isoflurane (3-5%) and then placed on a heated platform (FHC, Inc.) in a stereotaxic nose cone to maintain anesthesia. A large incision was placed over the animal’s skull, and the connective tissue and muscle surrounding the skull was removed using a fine scalpel blade. A modified headplate was attached using dental acrylic (Metabond) and secured to the skull. For cortical imaging, the skull was thinned with a dental drill until transparent, or removed entirely and covered with saline or ringers solution. After surgery, the isoflurane levels were dropped to ∼<1% for imaging and electrophysiology, the procedures for which were identical to those for the awake animal. The animal’s vitals (heart rate and respiratory rate) were constantly measured for tracking anesthesia depth.

#### Histology

Histological procedures were similar to those utilized previously (Borden et al., 2017), to validate ArcLight in S1 and/or opsin expression in VPm thalamus. Histological samples were prepared by perfusing the animal transcardially with phosphate buffered saline (PBS) followed by 4% paraformaldehyde. Brains were post-fixed overnight in 4% paraformaldehyde then transferred to PBS before sectioning. Thick sections were cut using a vibratome (100 μm, Leica, VTS 1000) and either directly mounted or saved for staining. See Supplemental Figure S1 for histological validation of expression of halorhodopsin in VPm and ArcLight expression in cortex.

#### Integrate & Fire or Burst (IFB) Modeling

To further explore the surprising finding of relatively invariant baseline firing rates during hyperpolarization of VPm thalamus, we utilized a biophysically inspired model of thalamic burst/tonic firing. Specifically, we suggest that this finding was a consequence of the burst mechanism, and not the result of possible confounds related to previously reported changes in reversal potential of chloride during prolonged periods of halorhodopsin activation (Raimondo et al., 2012). The Integrate and Fire or Burst (IFB) model was derived from previously published models of thalamic function from the LGN (Smith et al., 2000; Lesica and Stanley, 2004; Lesica et al., 2006). In order to simulate the experimental parameters and account for changes in thalamic activity, some additional terms and parameters were added and adjusted. Additionally, we generated ongoing activity using two methods, either injected current noise or synaptic events, with both showing the same results. The results shown here use the synaptic event model where IPSCs and EPSCs are modulated as fixed inputs. The model itself was written and analyzed using custom scripts in Matlab 2016a.

The model is based on modifications of the standard integrate and fire model representing the effects of integrated synaptic currents on membrane voltage:

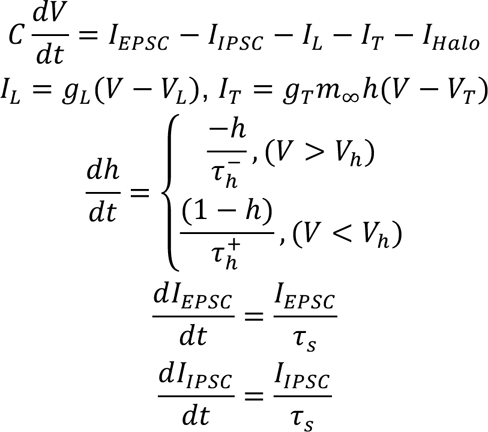

The following parameters were used to simulate thalamic activity: 𝐶 = 2uF/𝑐𝑚^2^, 𝑔*_L_* = 0.035 mS/𝑐𝑚^2^, 𝑔*_T_* = 0.07 mS/𝑐𝑚^2^, V*_L_* = −65mV, V*_reset_* = −45mV, V*_h_* = −68mV, V*_T_* = 120mV, 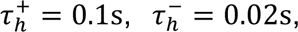 𝜏_0_ = 1e − 2uF/𝑐𝑚^2^, 𝑇ℎ𝑟𝑒𝑠ℎ𝑜𝑙𝑑 = −35mV, 𝐼*_Halo_* = 0 −1𝑢𝐴/𝑐𝑚^2^. EPSPs occurred at a rate EPSP*_rate_* = 1 − 25Hz, where each excitatory post- synaptic event had a peak of 3nA/𝑐𝑚^2^, and decayed according to the above first-order differential equation for I*_EPSC_*. Similarly, IPSPs occurred at a rate of IPSP*_rate_* = 0 − 5Hz, where each inhibitory post-synaptic event had a peak of 1𝑢𝐴/𝑐𝑚^2^, and decayed according to the above first-order differential equation for I*_IPSC_*. To simulate the different levels of thalamic activity, we varied the rates of EPSP inputs on the thalamic model (based on published ranges of thalamic activity (Urbain et al., 2015)). IPSPs were simulated at a much lower rate (20% of EPSP rate) to add additional variability to baseline activity. The model outputs represent the average response of 100 simulated thalamic neurons in response to various levels of thalamic hyperpolarization and baseline activity. The model was updated at 1ms steps. The absolute refractory period was set to 1ms.

#### Cortical E-I Modeling

We constructed a simple model of the thalamocortical network using custom scripts written in Python 3.6.10, as described previously (Wright et al., 2021) We modeled a single cortical barrel as a clustered network of excitatory and inhibitory single-compartment leaky integrate-and-fire (LIF) neurons, subject to excitatory thalamic and non-thalamic synaptic inputs. For each condition, we simulated 50 trials (200 ms per trial), with a time-step of 0.05 ms. A detailed description of the model can be found in Cortical E-I Modeling Supplemental, Section 1. All code is freely available upon request.

We created VPm input spiking to match both the experimentally observed sensory-evoked firing and the pair-wise synchrony of thalamic firing, across conditions. Specifically, we modeled a single VPm barreloid as forty independent “neurons”, or spike trains. Spike times were drawn from the empirical VPm PSTHs, re-scaled for each neuron to give a broad distribution of firing rates. Bursts were modeled as triplets of spikes with 1.25 ms inter-spike interval.

Non-zero thalamocortical (TC) synaptic weights were broadly distributed, and we implemented differential TC connectivity by imposing higher TC convergence (Bruno and Simons, 2002; Cruikshank et al., 2007) and shorter synaptic latencies (Cruikshank et al., 2007; Kimura et al., 2010) for inhibitory neurons, and requiring that VPm neurons with the highest mean rates synapsed only onto inhibitory neurons (Bruno and Simons, 2002).

We modeled a single cortical column as a network of 800 excitatory and 100 inhibitory LIF neurons, with relatively strong inhibitory-to-excitatory synapses (Gabernet et al., 2005). We imposed spatial clustering via “small-world” network connectivity (Litwin-Kumar and Doiron, 2012; Bujan et al., 2015; Wright et al., 2017b, 2017a), with 10% re-wiring probability. Inhibitory LIF neurons had shorter membrane time constants (Gentet et al., 2010) and refractory periods than excitatory neurons, which – together with the TC connection properties described above – supported higher firing rates in inhibitory neurons, as observed here (Fig. 7) and in previous work (Bruno and Simons, 2002; Khatri et al., 2004; Gentet et al., 2010; Taub et al., 2013). Excitatory neurons were subject to an inhibitory spike-rate adaptation conductance, which helped to stabilize network activity.

We employed alternate models to parse the roles played by changes in thalamic bursting per se, and by changes in synchronous thalamic spiking. In these alternate models, we manually manipulated tonic spike times to effect changes in the rate of synchronous tonic input spikes, while holding mean tonic and burst rates fixed. Specifically, for each trial and spike train, we probabilistically relocated tonic spikes that occurred within +/- 5 ms of the early PSTH peak to a later (random) time within approximately the first 30 ms post-stimulus.

For each model, we calculated the grand mean +/- SEM firing rates for all neurons, for an “early” (0 – 30 ms post-stimulus) and “late” (60 – 100 ms post-stimulus) response window, where stimulus onset time represents the time of galvo deflection onset, or *t = 0* in the empirical VPm PSTH. For cortical network synchrony analysis, we randomly selected 200 excitatory and 200 inhibitory neuron pairs. We then calculated synchrony for each “valid” pair, i.e., each of these pairs with at least one relative spike time in the 30 ms post-stimulus window across all trials. For VPm spike train synchrony analysis, we randomly selected 50 VPm neuron pairs, and repeated the procedure described above for the experimental VPm data. Note that we used this calculation to confirm that drawing thalamic spike times from the empirical VPm PSTHs, and assuming independence across these input spike trains, was sufficient to reproduce the empirical trends in synchronous thalamic spiking.

### Quantification and Statistical Analysis

#### Thalamic Electrophysiology Data Analysis - Mean Response, Burst Ratio, Synchronous Spike Counts

We report several different basic measurements of spiking activity from our thalamic units including evoked response and evoked bursting response. Thalamic firing activity was reported as PSTHs with units of firing rate in Hz, calculated as the number of spikes within a bin, divided by the size of the bin (see individual plots for bin size). We determined thalamic evoked response as the initial response (0-30ms) to sensory stimuli, reported as the number of spikes per stimulus averaged over many trials (9-102 trials). The corresponding evoked bursting response was determined as number of bursting spikes per trial in that same post stimulus period. Bursting spikes were defined as 2 or more spikes that fire at most 4ms apart preceded by 100ms of silence. The 100ms pre-stimulus activity is based on reported values for T-type calcium bursts (Lu et al., 1992; Swadlow and Gusev, 2001; Whitmire et al., 2016). All data analysis of the recorded extracellular thalamic units was accomplished using custom Matlab and Python scripts.

We calculated the mean synchronous spike counts using all pairs of sensory-responsive thalamic neurons recorded simultaneously using silicon probes, as described previously (Wright et al., 2021). Briefly, for each pair of simultaneously-recorded, sensory-responsive single-units, we considered the spike times in a brief (10 ms) post-stimulus window, and calculated the relative spike times (which ranged from +20 ms to -20 ms). We repeated this for all pairs, populating a grand set of relative spike times, separately for each light level, generating the spike cross-correlogram (CCG). We then summed the CCG between +5 ms and -5 ms, and divided by the number of contributing pairs. We repeated these steps using an equivalent number of random relative spike times, and subtracted the synchronous spike counts for these shuffled relative spike times from the true synchronous count. This yielded the “mean synchronous spike count (shuffle-corrected)”, or the average number of synchronous spike counts per pair, beyond what would be predicted from pairs of neurons with randomized relative spike times. We calculated 95% confidence intervals by re-sampling the relative spike times with replacement. We performed this analysis for three separate windows: 0 – 10 ms, 10 – 20 ms, and 20 – 30 ms post-stimulus.

#### Thalamic and Cortical Electrophysiology Data Analysis

Offline spike sorting was accomplished using either the Plexon Offline Spike Sorter v4 (Plexon, Inc, for tungsten electrode thalamic recordings), or KiloSort2, followed by curation in Phy (for silicon probe recordings in thalamus and cortex). We then required each unit to satisfy various criteria in order to be considered a well-isolated single-unit. We first calculated the signal-to-noise ratio (SNR), or the amplitude of the mean waveform (trough-to-peak) divided by the standard deviation. Second, we calculated the inter-spike-interval (ISI) violation percentage, or the percentage of all spikes within the 0 – 1 ms inter-spike interval. We required the SNR to be greater than or equal to 2 (3), and the ISI violation percentage to be less than or equal to 1 (1.5) for cortex (thalamus). Cortical single-units were classified as putative Regular Spiking Units (RSUs) or Fast Spiking Units (FSUs) based on the characteristics of the spike waveform (McCormick et al., 1985; Niell and Stryker, 2010; Guo et al., 2017; Speed et al., 2019; Yu et al., 2019). Specifically, units with a spike-width exceeding 0.4 ms (defined as trough-to-peak) were classified as RSUs, and below this as FSUs. For thalamic single-units, we required the spike-width to exceed 0.3 ms, as narrower waveforms could reflect activity at TRN synaptic terminals (Barthó et al., 2014). For multi-unit data, we measured threshold crossings from the continuously recorded thalamic or cortical activity. Thalamic multiunit activity was captured using a threshold criterion of 5 standard deviations over the entire recording (Yang et al., 2016). For cortical multiunit recordings, a manual threshold was set based on each experiment. Additional data analysis utilized custom scripts using Matlab (Mathworks, Inc).

Cortical units were initially analyzed together, across recording sites, as the dataset was not sufficient to confidently assess any cross-laminar effects. However, we did repeat some analyses using only putative layer 4 (L4) units, identified using methods similar to those described previously (Sederberg et al., 2019), as L4 was particularly important to this study as the primary thalamorecipient cortical layer. Briefly, for each experiment, we first calculated the across-trial average local field potential (LFP, or 2 – 200 Hz bandpass-filtered voltage traces) and multi-unit (MU, threshold crossings of 150 Hz highpass-filtered voltage traces) responses to punctate whisker deflections. We then calculated the current source density (CSD) profile of the LFP responses. Finally, we used a combination of the amplitudes and latencies of LFP deflections, the locations and latencies of CSD sinks, the amplitudes and latencies of MU firing, and the approximate probe depth to estimate the center of L4. Specifically, L4 is expected to be 400 – 600 µm below the cortical surface, and exhibit short-latency, large-amplitude LFP and MU responses, and a short-latency CSD sink. We required at least two of the three electrophysiological signals (LFP, CSD, MU) to provide qualitative agreement on the L4 center, and for this center to be between 400 and 600 µm below the estimated cortical surface, which excluded one of the six experiments from this analysis. Finally, we considered L4 channels to be those between 100 µm above and 100 µm below the central channel. A cortical single-unit was deemed a putative L4 unit if the channel on which its mean waveform was largest was included in the list of putative L4 channels.

Cortical firing activity was reported as PSTHs with units of firing rate in Hz, calculated as the number of spikes within a bin, divided by the size of the bin (see individual plots for bin size). All cortical spikes within a 1ms ISI were removed from analysis (and this was <1% of total). Synchrony across cortical single-units was computed from spike cross-correlograms across recorded pairs. Specifically, synchrony was defined as proportion of spikes from the full (+/- 100ms) cross-correlogram that were in a central +/- 7.5ms window (Wang et al., 2010; Whitmire et al., 2016). To determine the number of needed synchronous events to accurately measure synchrony, we simulated two neurons with a ∼5% change in synchrony (assuming a normal distribution). We found that approximately 50 events were required to accurately separate the two distributions.

#### Voltage Imaging Data Processing

Raw images were loaded and converted from the SciMedia “. gsd” format using custom scripts and down-sampled by a factor of two. Each dataset was first normalized to a %ΔF/F_o_ measurement by subtracting and dividing each trial by the temporal average of the frames 0 to 200ms preceding light delivery (F_o_). In two instances, where 200ms of preceding LED onset activity was not captured, the F_o_ was taken as an average 200ms period across no-stimulus trials. Hemodynamic noise was removed using a PCA background subtraction method. As described in detail (Borden et al., 2017), *in vivo* ArcLight imaging overlaps with the hemoglobin absorption spectrum, and therefore contains hemodynamic noise that must be removed for analysis. Imaging the wildtype mouse cortical surface using the same blue excitation and ArcLight filter set revealed similar patterns of oscillatory activity, likely through auto-fluorescence and effects of hemodynamic absorption and blood flow (Ma et al., 2016). The background PCA subtraction utilizes the auto-fluorescence signal from non-ArcLight transfected regions to predict the hemodynamic signal across the recorded space. Specifically, the method uses principal component analysis of non-expressing low background auto-fluorescence regions (determined from the maximum fluorescence from a non-injected animal) to find the ongoing hemodynamic components on a single trial basis. Additionally, the background fluorescence regions were selected at least 1mm away from the recorded whisker evoked response (Borden et al., 2017). Ideally, these criteria would create a spatially defined region with little or no ArcLight fluorescence to isolate the hemodynamic signal from the signal of interest. Each frame is first spatially averaged by either a 200 µm x 200 µm circular averaging (pillbox) filter or a media filter to reduce noise. On a single trial, the corresponding top five principal components of the low background regions (which contains approximately 85% of the variance explained) are projected on a pixel by pixel basis across the entire recording using a lasso regression method with regularization. The lasso regression utilizes a cross-validated approach to determine the minimum number of components to develop the model of hemodynamic noise. In order to prevent the removal of any stimulus evoked activity, each pixel was fit on pre-stimulus activity (either before light onset for experiments involving optogenetics, or immediately preceding stimulus delivery). The final predicted hemodynamic signal for each pixel was subtracted across the entire recording on a pixel by pixel basis. Due to the complex waveform of the hemodynamic response, a simple notch filter is not effective at separating the signal from the noise (Borden et al., 2017). We found that the background PCA subtraction method greatly reduced hemodynamic signal across the entire frame, compared to the off-ROI method (Borden et al., 2017). In some instances, brief onset and offset light artifacts of the 590nm light was visible in the recorded ArcLight cortical signal. To account for any optogenetic transient light artifacts, we only considered the relative changes in fluorescence during steady state light levels. Both raw and processed images showed qualitatively similar results.

#### Awake Voltage Imaging Data Analysis – Dual Camera

In the awake animal, we utilized a dual camera imaging system to capture a background fluorescence signal for hemodynamic subtraction. Two different cameras were used to capture the ArcLight and auto-fluorescence signals, and therefore, pixels could not be directly registered for subtraction for pixel by pixel correction. Instead, we utilized the same Background PCA subtraction method to find and develop models of the hemodynamic response based on the global PCA signal derived from the background image. For the dual camera data, each component was fit over the entire recording for subtraction of the hemodynamic noise. Both raw and processed images showed qualitatively similar results. Unless otherwise noted, each dataset was processed with the Background PCA or Dual Camera subtraction method as stated above.

#### Imaging Data Analysis – Peak amplitude, Normalized Peak, and Temporal Properties

We measured the effect of the optogenetic stimulation on the peak amplitude of the evoked mean ArcLight fluorescence in the determined cortical barrel. The cortical barrel region of interest (ROI) for each stimulated barrel and each data set was selected as the ∼200 µm x 200 µm region with the largest response 30ms after stimulus delivery. This determined ROI was used for all subsequent analyses of the temporal response. To better isolate the evoked amplitude, the frame preceding stimulus delivery (t=-5ms) was subtracted from the resulting recorded signal. For each recording, the peak amplitude was defined as the ΔF/F_o_ at the time of the maximum average response between 0 and 110ms for the strongest stimuli (1200 Deg/s) presented under control and various optogenetic conditions. In order to measure the temporal properties of the evoked response, we concentrated on the timeseries data from the determined cortical barrel ROI. For normalized fluorescence (Norm ΔF/ F_o_), each session’s peak response was divided by the average peak response to the strongest stimulus (1200 Deg/s) under the control condition. The normalization allows for a better comparison across animals which may have different levels of ArcLight expression. Peak time was defined as the time of the maximum fluorescence between 0 and 110 ms post stimulus, and the time of return to baseline as the time when the fluorescence crossed the pre-stimulus baseline value following the peak. Peak-to-baseline was then calculated as the time between the fluorescence peak and the return of the fluorescence to baseline. Recovery was defined as the average fluorescence in the 120-400ms window following the stimulus.

#### Imaging Data Analysis – Area Measurements

In addition to measuring the peak response, we also measured the effect of different thalamic states on the evoked area of sensory cortical activity. We measured the activated area as the number of pixels exceeding a threshold using the average response at the peak frame (0-110ms) preceding stimulus delivery. Similar to other studies (Lustig et al., 2013; Millard et al., 2015), we measured the spatial activation using the 70% threshold. To compare the area independent of amplitude changes, we normalized the peak frame by dividing by the peak fluorescence in each condition (Control and LED). In order to isolate the evoked activity from ongoing activity, we subtracted the frame preceding stimulus delivery (t=-5ms). Different thresholds had no effect on the observed trends.

#### Statistical Analysis

All tests were conducted using the MATLAB Statistics Toolbox (Mathworks, Inc.). For all measurements, we determined if the specific data sets were normally distributed using the Lilliefors test for normality. If the data were normal, we used the appropriate (paired or unpaired) t-test for statistical difference. If the population was determined to have non-normal distributions, we conducted non-parametric Wilcoxon signed-rank tests to determine statistical significance. All sample sizes are reported in the figure captions and Results text, along with an indication of particular test and the corresponding statistical significance level (* - p<0.05, ** - p<0.01, *** - p<0.001 in figures).

