## Supplemental Information for "Thalamic bursting and the role of timing and synchrony in thalamocortical signaling in the awake mouse"

#### Figure S1

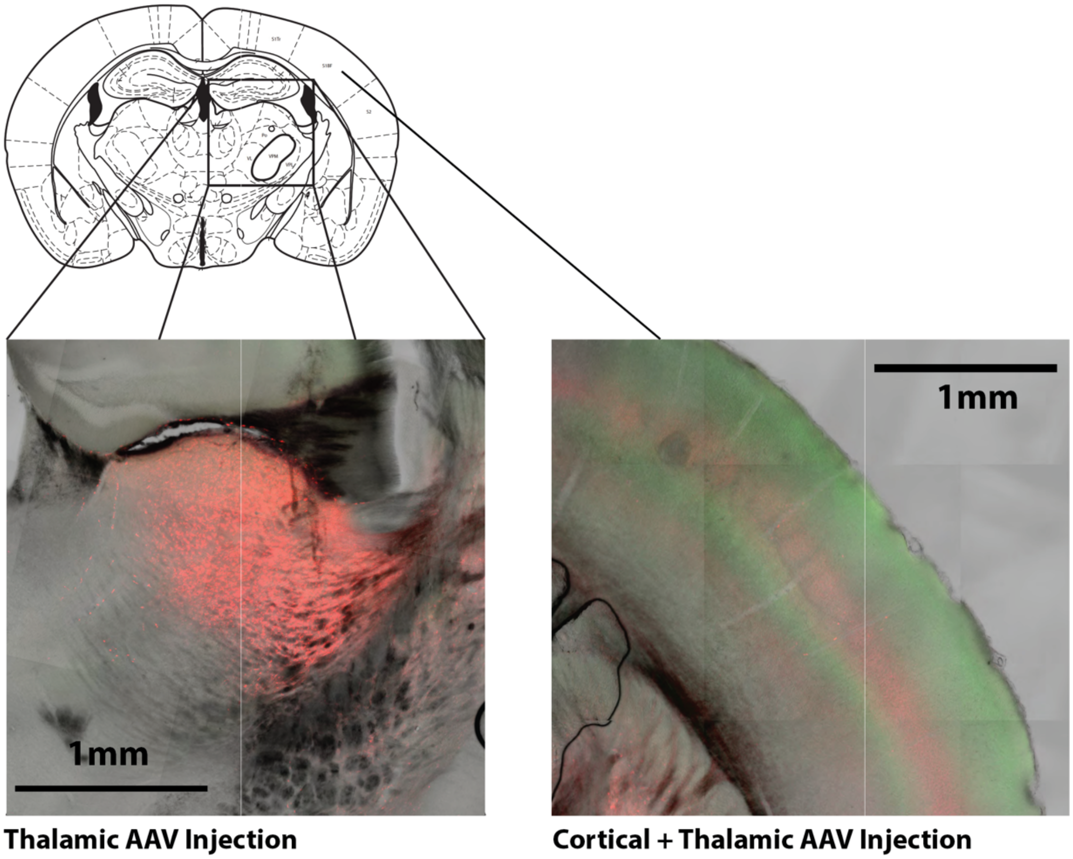


**Supplemental Figure S1. Opsin and GEVI controls. Example of combined Halorhodopsin and ArcLight expression in Mouse Sections. Left.** Expression of Halorhodopsin in the mouse thalamus. Mice are injected with two viral vectors, ArcLight in cortex (AAV1-hysn1- ArcLightD-SV40), and eNphR3.0 (AAV5-CamIIKianse-eNphR3.0-mCherry) in VPm thalamus. Thalamic expression of Halorhodopsin (mcherry-red- Emission 608-715nm) is localized throughout the thalamic region of the mouse (red). Electrode tracks of the optrode (optic fiber and electrode) are seen terminating in the VPm region. **Right.** Cortical injection of ArcLight probe reveals expression throughout layer 2/3 and layer 5 across the mouse cortex (ArcLight-green-Emission 474-562nm). Thalamic expression of Halorhodopsin (mcherry-red) is also found in the axons of the thalamic neurons projecting to layer 4 and 5 of the S1 barrel cortex.

### Figure S2

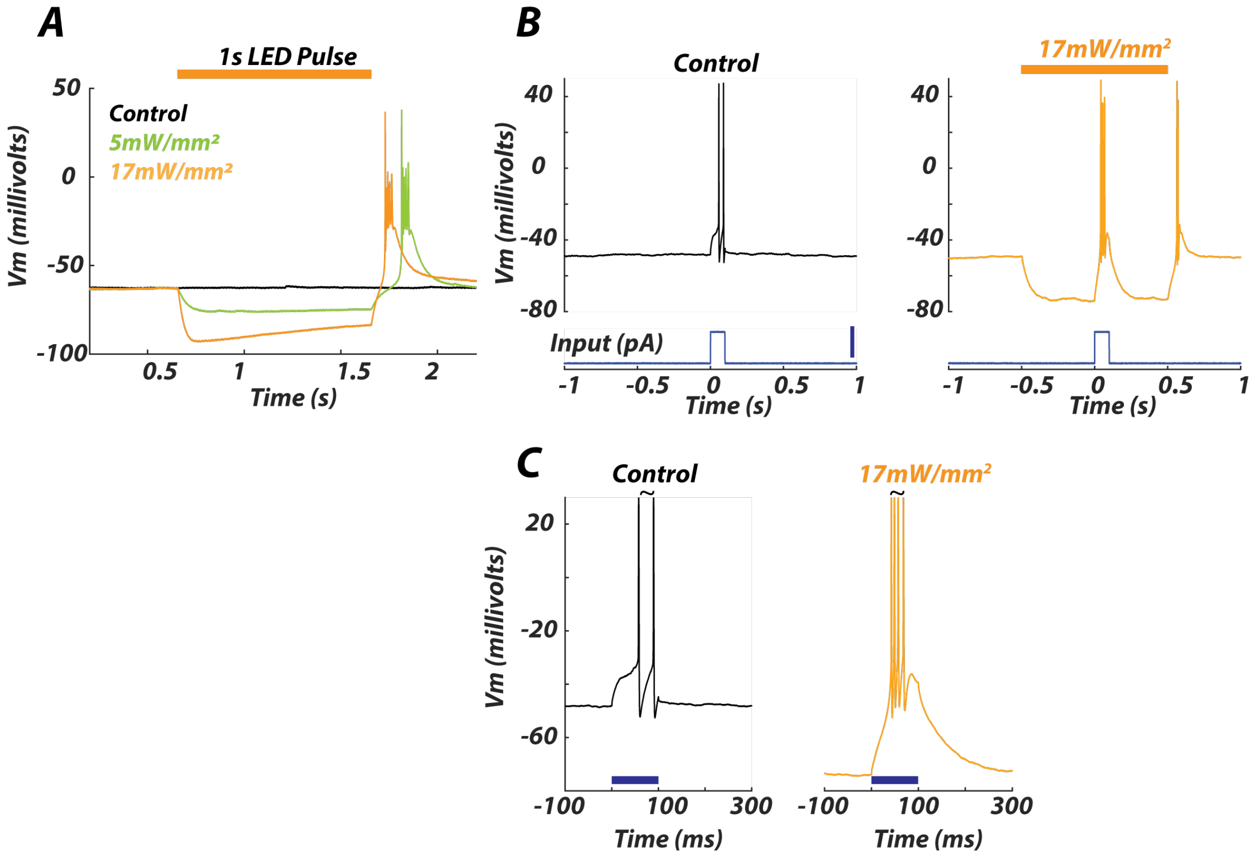


**Figure S2. Intracellular brain slice recordings from thalamic neurons transfected with halorhodopsin show burst behavior after periods of hyperpolarization.** Methods were taken from Zhao et al., 2011. Mice were anesthetized and perfused with 25–30 mL of carbogenated protective artificial cerebrospinal fluid (aCSF) of the following composition: 92 mM N-methyl-D-glucamine (NMDG), 2.5 mM KCl, 1.25 mM $NaH_2PO_4$, 30 mM $NaHCO_3$, 20 mM HEPES, 25 mM glucose, 2 mM thiourea, 5 mM Na-ascorbate, 3 mM Na-pyruvate, 0.5 mM $CaCl_2.4H_2O$, and 10 mM $MgSO_4.7H_2O$. The pH of the solution was titrated to 7.3–7.4 with concentrated HCl. Brains were embedded in 2% agarose and mounted for coronal sections 300 μm thickness. Slices were recovered for ≤ 20–30 minutes at room temperature (23–25 °C) in carbogenated protective cutting aCSF. After this initial recovery period the slices were transferred into a chamber containing room temperature carbogenated aCSF of the following composition: 119 mM NaCl, 2.5 mM KCl, 1.25 mM $NaH_2PO_4$, 26 mM $NaHCO_3$, 12.5 mM glucose, 2 mM $CaCl_2.4H_2O$, 2 mM $MgSO_4.7H_2O$. The aCSF was supplemented with 2 mM thiourea, 5 mM Na-ascorbate, and 3 mM Na-pyruvate, and slices were stored for 1–5 hours prior to transfer to the recording chamber for use. The osmolarity of all solutions was measured at 300–310 mOsm and the pH was maintained at ~7.3 after equilibration under constant carbogenation. The slices were perfused with room temperature (22–25 °C) carbogenated recording aCSF at a rate of 4 mL per min. Whole-cell patch-clamp recordings were obtained from visually identified neurons using boroscilicate glass pipettes pulled on a horizontal pipette puller (Sutter Instruments) to a resistance of 3–8 MΩ when filled with the internal solution containing 145 mM K-Gluconate, 10 mM HEPES, 1 mM EGTA, 2 mM Mg-ATP, 0.3 mM $Na^2$-GTP, and 2 mM MgCl. Neurons expressing Halorhodopsin were identified by visualization of membrane-targeted mCherry or YFP fluorescence. Amber laser light (590 nm) was delivered through a 200 μm diameter optic fiber (ThorLabs) positioned near the recorded neuron. The other end of the optic fiber was coupled to an LED light source (ThorLabs). Current pulses were delivered in current clamp using Clampex software. **A.** 1s LED step inputs applied during whole cell recording of thalamic neuron with increasing levels of LED input. Halorhodopsin activation induces a hyperpolarizing current into the thalamic neuron (Control- Black, approx. 5mW/mW$^2$ - Green, and approx. 17mW/mm$^2$ – Orange). After the cessation of optogenetic input, the thalamic neuron responds with a characteristic post-inhibitory rebound due to T-type calcium channel de-inactivation (1 Trial). **B.** Ongoing polarization alters the encoding of the same step current input into thalamic neuron. Under control conditions (**Left**), the thalamic cell responded to a depolarizing current step with a tonic firing of two action potentials. After 500ms of hyperpolarizing Halo activation (17mW/mm$^2$), the thalamic cell produces a burst response to the same current step. Blue bar represents 100pA stimulation. **C.** Thalamic polarization can modulate the evoked response to current inputs. Same input as shown in B, however across three different light levels of Halo activation (**Left:** Control – Black; **Right:** 17mW/mm2 - Orange). Blue bar represents duration of 100pA stimulation. Note, action potentials marked with (~) extend beyond axis.

### Figure S3

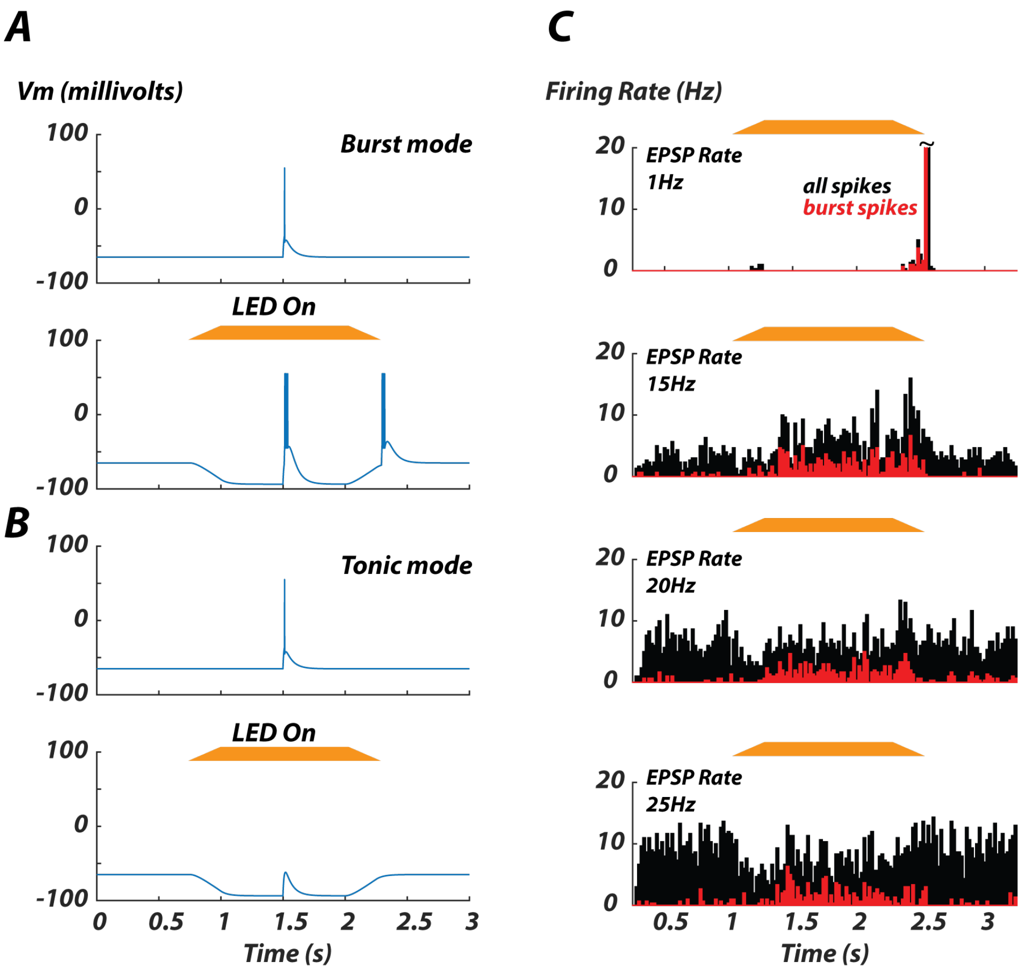


**Supplemental Figure S3. Integrate and Fire & Burst (IF&B) model neuron replicates in vivo responses to Halorhodopsin Input.** While prolonged halorhodopsin stimulation has shown to change the reversal potential for chloride (Raimondo et al., 2012) we show through simulation here that T-type calcium channels found in thalamic neurons are sufficient for producing the effect we observe. The simulated neuron with a simple low voltage burst mode replicated what we observed in the awake (high background activity, Figure 2) and anesthetized animal (low background activity, Figure 1). **A.** IF&B model neuron replicates thalamic bursting and post-inhibitory rebound **A. Top** - Control strong current injection evoked a single tonic spike at t=1.5s. **A. Bottom** - Thalamic hyperpolarization (through simulated halorhodopsin current) causes thalamic burst to same current input as top, and a post-inhibitory rebound event. **B.** Same inputs as in A, however IF&B model does not contain fictive T-type calcium channels. No bursting event occurs during hyperpolarization. For A-B (with no spontaneous inputs), we increased the current injection 3x to simulate a strong input. **C.** IF&B model PSTH responses (100 trials) to various spontaneous activity rates (from 1-25Hz simulated Excitatory Post Synaptic Potentials (EPSPs)) during an ongoing hyperpolarizing input with all spikes (black) and burst spikes (red). Note, as the ongoing input increases, a hyperpolarizing input changes the net effect (from increasing ongoing activity to replacing ongoing activity with bursting events). Note, simulated firing rate marked with (~) extend beyond axis.

### Figure S4

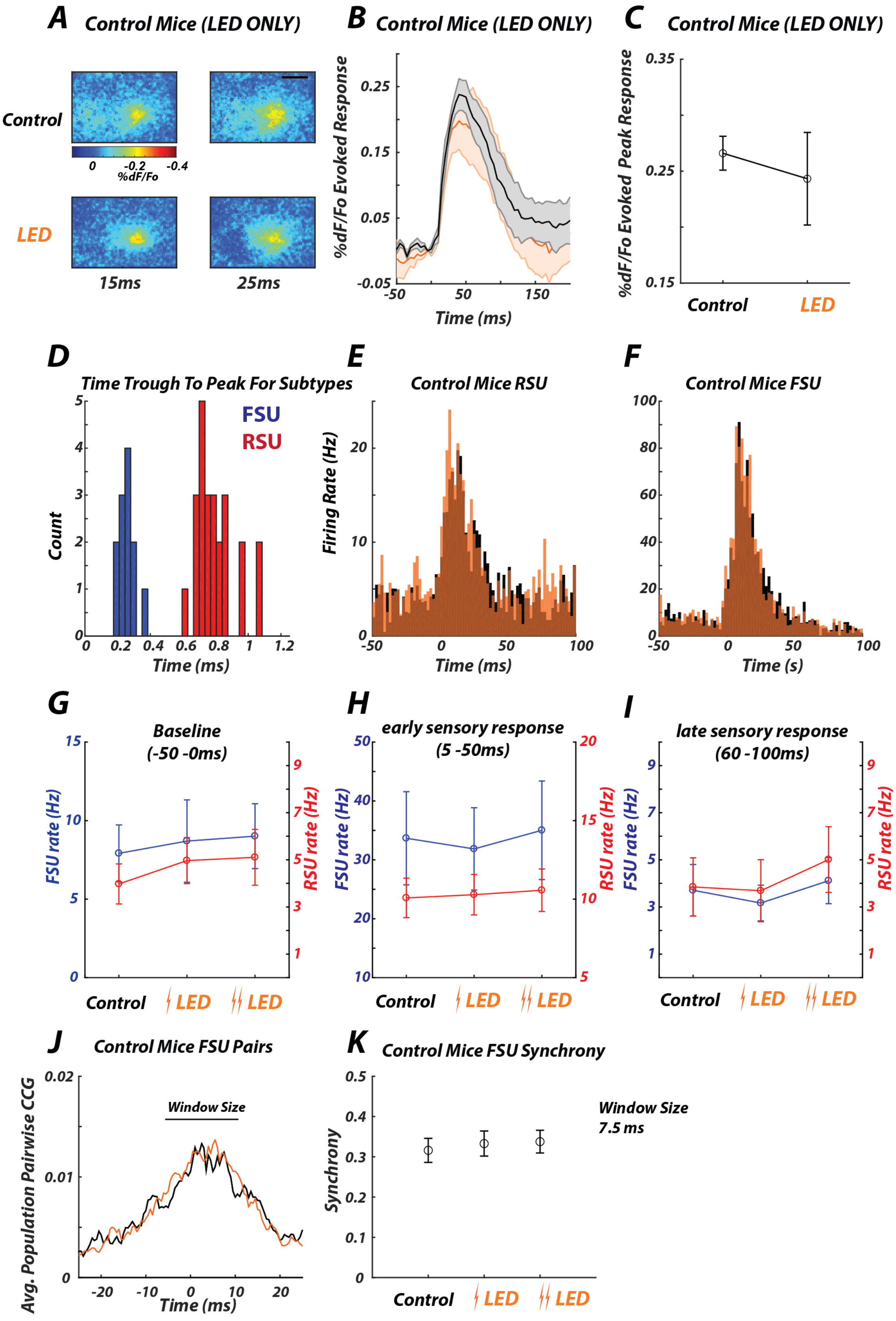

A Control Mice (LED ONLY)
Control
LED
0
-0.2
-0.4
%dF/Fo
15ms
25ms
B Control Mice (LED ONLY)
%dF/Fo Evoked Response
Time (ms)
C Control Mice (LED ONLY)
%dF/Fo Evoked Peak Response
Control
LED
D
Time Trough To Peak For Subtypes
FSU
RSU
Count
Time (ms)
E
Control Mice RSU
Firing Rate (Hz)
Time (ms)
F
Control Mice FSU
Time (s)
G
Baseline
(-50 -0ms)
FSU rate (Hz)
RSU rate (Hz)
Control
LED
LED
H
early sensory response
(5 -50ms)
FSU rate (Hz)
RSU rate (Hz)
Control
LED
LED
I
late sensory response
(60 -100ms)
FSU rate (Hz)
RSU rate (Hz)
Control
LED
LED
J
Control Mice FSU Pairs
Avg. Population Pairwise CCG
Window Size
Time (ms)
K
Control Mice FSU Synchrony
Synchrony
Window Size
7.5 ms
Control
LED
LED

**Supplemental Figure S4. Control for impacts of LED in WT Mice, non-injected with halorhodopsin. A.** Example of average evoked GEVI spatial response (51 trials) in non-halorhodopsin injected mice across no light (control) and LED on conditions (17mW/mm$^2$)**.** Black bar represents 1mm. **B.** Average timeseries response in the evoked ROI (see methods) across no light control (Black) and LED On (Orange) conditions across 6 recording sessions (2 mice) **C.** Peak Evoked GEVI response across all animals (n=6 recording sessions, 2 mice) We observed no trend in the evoked response due to the LED. **D.** Breakdown of Cortical fast spiking units (FSU) (n=13) and regular spiking units (RSU) (n=26) in non-halorhodopin injected mice. Units classified by the trough-to-peak time (<0.4ms, see methods). **E-I.** Impact of LED on cortical evoked response in non-halorhodopsin injected mice. We observed no impact on the dynamics of the evoked response (bin size=2ms). **J.** Cross correlation across FSU pairs (n=13 pairs). Cross-correlograms smoothed via a moving average filter, 2.5ms window. **K.** Synchrony metric in non-injected mice across FSU pairs (see Figure 7). Whisker stimulus delivered at time 0. Error bars represent mean +/- SEM.

### Figure S5

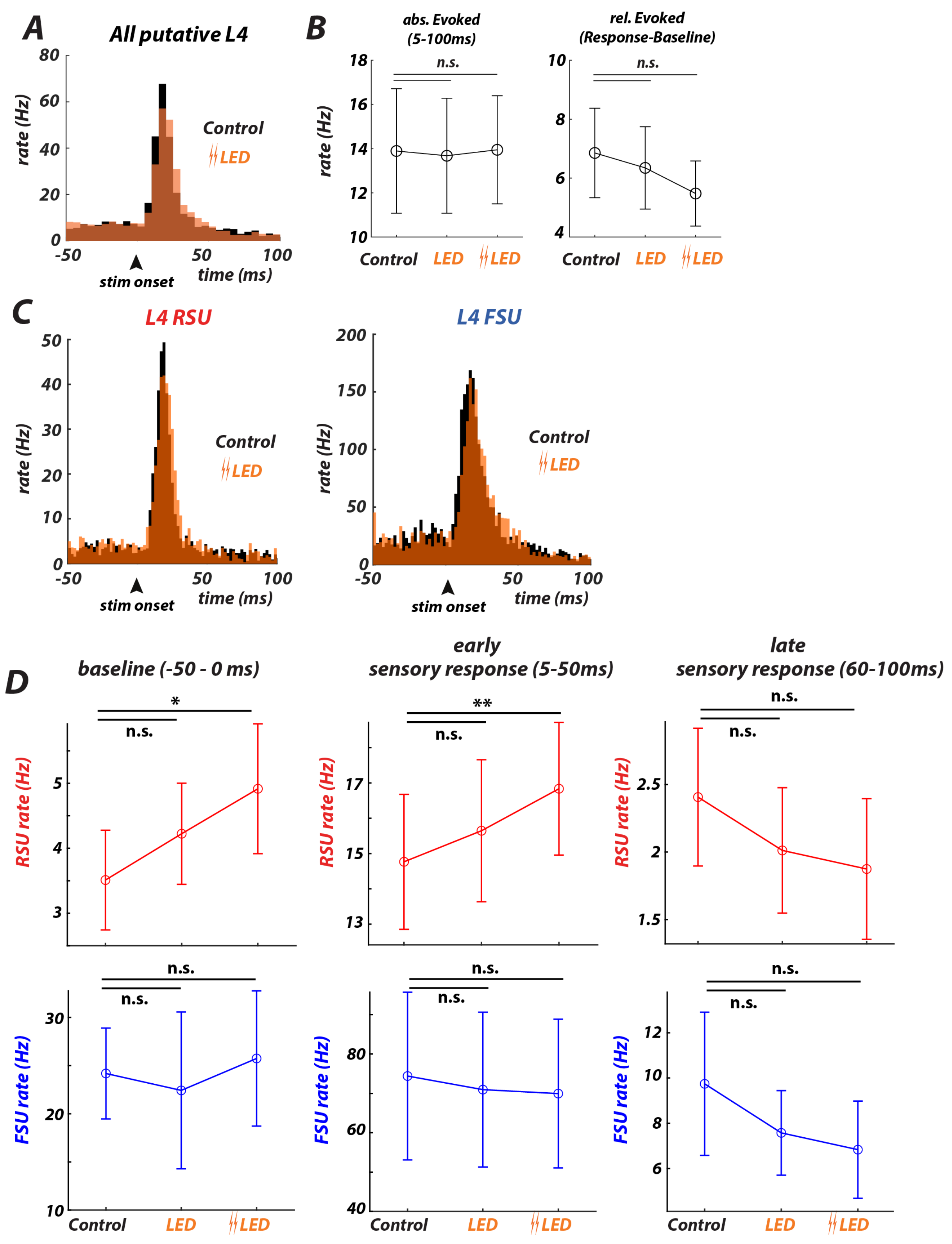

A
All putative L4
rate (Hz)
80
60
40
20
0
-50
50
100
stim onset
time (ms)
Control
LED
B
abs. Evoked
(5-100ms)
rel. Evoked
(Response-Baseline)
rate (Hz)
18
16
14
12
10
n.s.
8
6
4
Control
LED
LED
C
L4 RSU
L4 FSU
rate (Hz)
50
40
30
20
10
200
150
100
Control
LED
stim onset
time (ms)
D
baseline (-50 - 0 ms)
early
sensory response (5-50ms)
late
sensory response (60-100ms)
*
**
n.s.
RSU rate (Hz)
5
4
3
17
15
13
2.5
2
1.5
FSU rate (Hz)
30
20
10
80
60
40
12
8
6
Control
LED
LED

**Supplemental Figure S5. Thalamic hyperpolarization affects timing but not magnitude of sensory-evoked responses in cortical L4. A.** Cortical Grand PSTH evoked sensory response (t=0) across all recorded single units (n=41 units, 5ms bins) for Control (black) and LED (orange) conditions. **B. Left.** Average absolute evoked cortical response remained approximately invariant during LED on conditions across all cortical recorded units (Control vs LED p=0.9, Control vs Double LED p=0.5, paired Wilcoxon signrank, n=41 units). Evoked sensory response period defined as between 5-100ms post stimulus. **Right.** Relative evoked cortical response exhibited a decreasing but not significant trend with increasing LED intensities (Control vs LED p=0.6, Control vs Double LED p=0.6, paired Wilcoxon signrank, n=41 units). Relative evoked response defined as the absolute response minus the preceding baseline activity -50-0ms pre-stimulus. Error bars represent mean +/- SEM. **C.** PSTHs of the aggregate putative RSUs (left, n=34) and FSUs (right, n=7) in response to a punctate whisker deflection at time t=0 (bin size 2ms) for the Control (black) and double LED (orange) conditions. **D.** Summary analyses (Error bars represent mean +/- SEM) for RSUs (top row) and FSUs (bottom row). Left - Mean baseline (-50 to 0ms) RSU (Control vs LED p=0.1, Control vs Double LED p=0.046, Wilcoxon signed-rank test, n=34) and FSU (Control vs LED p=0.3, Control vs Double LED p=0.6, Wilcoxon signed-rank test, n=7) firing rates for the LED relative to the Control condition. Middle - RSU (Control vs LED p=0.3, Control vs Double LED p=0.046, Wilcoxon signed-rank test, n=34) and FSU (Control vs LED, p=0.6, Control vs Double LED p=0.6, Wilcoxon signed-rank test, n=7) early sensory response (5-50ms) for the LED compared to the Control conditions. Right – Late sensory response (60-100ms) for the LED compared to the Control condition for the RSU (Control vs LED p=0.3, Control vs Double LED p=0.15, n=34, Wilcoxon signed-rank test) and FSU (Control vs LED p=0.3, Control vs Double LED p=0.6, n=7, Wilcoxon signed-rank test) populations. LED approx. 17mW/mm$^2$ ramp, Double LED approx. 35mW/mm$^2$ ramp – see Methods. Error bars represent mean +/- SEM.

#### Supplemental Information on Cortical E-I Modeling

We constructed a simple model of the thalamocortical network using custom scripts written in Python 3.6.10, using the same model parameters as described in detail previously (Wright et al. 2021), except where noted below. All code is freely available upon request. We modeled a single cortical barrel as a clustered network of excitatory and inhibitory single-compartment leaky integrate-and-fire neurons, subject to thalamic and non-thalamic excitatory synaptic inputs. We selected numbers of model VPm and S1 neurons, and thalamocortical and intracortical connection parameters such that i) there was a numerical expansion of neurons at the thalamocortical junction; ii) the majority of synaptic inputs to a cortical neuron arose from other cortical neurons; and iii) cortical excitatory neurons were subject to strong feedforward inhibition. As such, the activity of the cortical network was not a simple read-out of thalamic spiking.

***Model of VPm Barreloid.*** We modeled a single VPm barreloid as forty independent spike trains. The grand mean pre-stimulus firing rate was set equal to the empirical grand-mean VPm rate in a 30 ms pre-stimulus window for each condition (9.14 Hz for control and 7.34 Hz for LED, calculated from the 46 sensory-responsive VPm units recorded via silicon probe). The ongoing and evoked rates for each neuron were then multiplied by a rate modulation factor drawn from a skewed gamma distribution (with a shape value of 2.0, a scale value of 1.0, then re-scaled to have a mean value of 1.0), to mimic the broad firing rate distributions of VPm neurons previously reported (Pinto et al., 2000; Bruno and Simons, 2002; Wang et al., 2010; Whitmire et al., 2016). Tonic and burst spike times for a given neuron were then drawn from their associated empirical PSTHs, multiplied by the neuron's rate factor. For tonic spikes, we required a minimum inter-spike interval (ISI) of 4 ms for all spikes drawn for a given neuron and trial. We modeled bursts as triplets of spikes: we first drew one spike time from the empirical burst PSTH, and then added spikes 1.25 ms before and after this central spike. We further required at least 2 ms of silence before the first spike in each burst, and excluded any bursts with spike times that fell outside of the trial window. Model VPm neurons were independent, in that spike times for each neuron were drawn independently. In this standard model, we did not make any additional assumptions regarding evoked VPm synchrony in the control and LED conditions, but by construction this resulted in across-condition synchrony effects that were consistent with the experimental observations. In our alternate models, we manually manipulated the rate of synchronous tonic spiking across VPm spike trains, while holding the rate of tonic and burst spikes fixed. Specifically, for each sensory-evoked tonic VPm spike that occurred within +/- 5 ms of the VPm tonic PSTH peak, we moved the spike away from the PSTH peak with probability P (repeating this simulation for P = 0.05, 0.15, 0.25). The new spike time was the grand PSTH peak time plus a random value. To obtain this random value, we first drew a spike time from a normal distribution with zero mean and standard deviation of 20 ms, took the absolute value, and added 5 ms.

***Model architecture and intrinsic neuronal properties.*** Each VPm neuron synapsed onto a subset of cortical network neurons via a set of thalamocortical (TC) synapses, with at most one synapse between each VPm and each network neuron. As done previously (Wright et al. 2021), we implemented differential TC connectivity by setting higher TC synaptic convergence (Bruno and Simons, 2002; Cruikshank et al., 2007) and shorter synaptic latencies (Cruikshank et al., 2007; Kimura et al., 2010) for inhibitory than for excitatory neurons, and also by requiring that VPm neurons with the highest firing rates synapse exclusively onto inhibitory neurons (Bruno and Simons, 2002), which generally supported strong feedforward inhibition in this model network.

Each thalamic spike resulted in a postsynaptic conductance in each postsynaptic cortical neuron, and we tuned the TC synaptic conductance amplitude such that near-simultaneous firing of multiple thalamic neurons was required to evoke action potentials in target neurons (Bruno and Sakmann, 2006) when intracortical synaptic strengths were set to zero.

We modeled a single cortical column as a network of 800 excitatory and 100 inhibitory neurons with “small-world” connectivity, as described previously (Wright et al. 2021). We tuned network connectivity parameters such that thalamic and random external inputs together could evoke bouts of persistent network firing on a given trial, but without runaway firing (see Table S7).

For network LIF neurons, we selected intrinsic excitatory and inhibitory neuronal properties that were consistent with previous modeling studies, and/or were motivated by previous experimental work, as described previously (Wright et al. 2021). Generally, inhibitory neurons had shorter membrane time constants and refractory periods than excitatory neurons, and excitatory neurons were subject to spike-rate adaptation.

For each condition, we simulated 50 trials, each lasting 200 ms (including a 50 ms “buffer window” to allow the network to reach stead-state, a 50 ms “pre-stimulus” window, and a 100 ms “post-stimulus” window), with a time-step of 0.05 ms. At each time-step, the membrane potential V of a given network neuron evolved according to its synaptic inputs, as described previously (Wright et al. 2021).

***Comparison to empirical S1 spiking responses.*** When tuning model parameters to reproduce the empirical results of interest, we constrained the model to qualitatively reproduce a relatively fast rise and slow decay in S1 firing rate after stimulus onset, with higher inhibitory than excitatory firing rates, while avoiding network saturation (i.e., individual neurons firing at the maximum rate allowed by the imposed refractory periods, which was 500 Hz for excitatory neurons, and 1000 Hz for inhibitory neurons). Given the simplicity of this model relative to the mammalian thalamocortical network, we did not attempt to reproduce the exact time-course of the empirical sensory response, nor the empirical firing rates of both excitatory and inhibitory neurons. Of note, the ratio of inhibitory to excitatory firing rates in the model was much higher than in the experimental data, and evoked spiking in the model returned to baseline faster than we observed in experiment. It is likely that adding additional features to the model could reproduce these results. For example, we modeled a single population of fast-spiking (PV) inhibitory neurons, but adding a second population of inhibitory neurons that functionally mimics the L4 SOM network would tend to inhibit PV interneurons (Ma et al., 2012; Xu et al., 2013) and thus disinhibit excitatory firing (Xu et al., 2013), particularly later in the sensory response, as these neurons have higher response latencies (Yu et al., 2019), and are driven by facilitating synapses (Beierlein et al., 2003; Hu and Agmon, 2016). Because our goal was to instead to reproduce our central experimental results with the simplest possible model, and because we could not segregate our recorded inhibitory units into PV and SOM interneurons with the experimental methods employed here, we did not pursue this goal in the current study.
